# Monkeys use the rod-dense retinal region rather than the fovea to visually fixate small targets in scotopic vision

**DOI:** 10.1101/290759

**Authors:** Oleg Spivak, Peter Thier, Shabtai Barash

## Abstract

Monkeys appear to visually fixate targets in scotopic conditions. The function fixations fulfill in photopic vision, keeping the target’s image on the fovea, is nullified in scotopic vision, because the fovea, with its cones, is desensitized in dim light. Here we followed the hypothesis that a previously described retinal region, the locus of maximal rod density, functionally replaces the fovea; we found that with dark background, most of the fixations direct the fovea above the target, so that the target’s image appears to fall on the line connecting the fovea with the locus of maximal rod density. There is considerable trial-by-trial variation in the fixation positions along this line. On the whole, the closer the visual conditions are to full scotopic, the higher is this gaze upshift, indicating the closer does the target fall to the locus of maximal rod density. Mesopic background induces low mean upshift. Full (45-min) dark adaptation was essential to achieving high upshift values. There is no analogous photopic effect – 45-min ‘bright adaptation’ did not shift the locus of photopic fixation.

## Introduction

### Fixation and saccadic eye movements are crucial for photopic vision

The evolution of a distinct, well-defined fovea is thought to be a signature of the primate and human visual system. The fovea comprises a much higher density of cones than the periphery; its boundaries are sharp; it is well defined anatomically and physiologically; and the specialized processing of foveal information goes far beyond the receptor level, through the retina’s other structure and into the brain. In visual cortex, information from the fovea is vastly over-represented. The high receptor density, couples with the special circuitry spanning retina through visual cortex, underlie the fovea’s unique capacity for high-acuity vision.

Nonetheless, the fovea is small. At any one time, the fovea covers only a small patch of the visual scene. To allow for high-acuity vision of more than a small patch, the eye must be rotated in the orbit, so that the fovea covers the next location of interest. Thus specialized eye movements have evolved in parallel to the evolution of the fovea. These movements stabilize the eyes during brief ‘fixations’, which allow vision to take place; and then shift the eyes, briskly yet precisely, to the next location (‘saccades’). Fixation and saccade eye movements are crucial for vision.

### The seeming paradox of fixation and saccades in scotopic vision

In dim light, cones are desensitized. Because most of the fovea is made up of cones, in scotopic vision the fovea is largely blind. The rationale for the function of saccades breaks down. Why keep shifting the target’s image to the fovea, now that the fovea’s visual performance is inferior to the rest of the retina? Nonetheless, seemingly paradoxically, the pattern of fixations and saccades is sustained in scotopic vision. Why?

### Rod-dense region in the superior retina

It had been known for a long time that rod density is not uniform over the retina, varying at least with eccentricity: from no rods at the center of the fovea, rod density increases with eccentricity to a peak at about the eccentricity of the blind spot. This peak reflects the ‘rod ring’, a circle of high rod density centered on the fovea. At higher eccentricities, rod density subsides, reflecting the ubiquity of very large rods in the far periphery.

With the advent of computerized histological technique, cone and rod density could be measured in the 2-dimensional manifold of the retina. As expected, cone density had a single peak, in the fovea, with a value much higher than cone density in the periphery. Rod density was found to have a more complex distribution. Studies by the groups of Rakic (Wikler and Rakic, 1990; Wikler et al., 1990) and of Curcio (Packer et al., 1989; Curcio and Allen, 1990) found a region of high rod density in the superior retina. Generally, the superior retina was found to have more rods than the inferior retina. The rod-dense region was called ‘rod hotspot’ or ‘dorsal rod peak’. Not wanting to commit to the precise details of the anatomy, in the present paper we refer to a rod-dense region, with the idea that it corresponds at least roughly to the region described in these studies.

The rod-dense region is well defined, but it does not have the crystal-clear structure of the fovea. The boundaries of the fovea are sharp; those of the rod-dense region, much less.

### Hypothesis: in scotopic vision, the rod-dense region might replace the fovea

Here we follow the hypothesis that the rod-dense region has a distinct function in scotopic vision, which is a variation of high-acuity vision. Thus, the function of the rod-dense region in scotopic vision is not unlike that of the fovea in photopic vision. We speculate that beyond the general similarity there might be important differences between the functions of the fovea and the rod-dense region, since photopic and scotopic vision evidently are not the same.

The hypothesis leads to a testable prediction: in scotopic vision, the image of the target would be fixated by the rod-dense region, or, at least, by its vicinity; not by the fovea.

This hypothesis offers a possible solution to the paradox of why the pattern of fixations goes on in the dark, even though the fovea is desensitized. Indeed, if the hypothesis would hold, it is not the fovea that is used in scotopic vision – but another retinal region, that in scotopic vision might enable relatively high acuity.

The hypothesis was first suggested in 1998 (Barash et al., 1998) but has not been tested to date. To understand why, we need to recall the time-course of dark adaptation.

### Distinction between photopic and scotopic dark

The time-course of dark adaptation is marked by two phases (Normann and Werblin, 1974). While staying in dark, a subject’s visual sensitivity increases. The time-course of the sensitivity has two exponential phases. On going from a bright environment to dark, the first exponential phase, lasting typically about 15 min, reflects the adaptation of the cones. At this time the rods are still saturated by the bright environment that preceded the dark. Only after the cone-based sensitivity stabilizes does the rod sensitivity start to dominate vision. It takes about 45 min for the process to near completion. (Because the rod-based sensitivity also increases exponentially, the time of completion is not well defined, but 45 min appears to be a conservative figure).

The perception of stimuli appearing during the first and second phases of the dark-adaptation time-course is likely to differ, because the first phase involves primarily cones, and the second primarily rods. Therefore, we call the first phase ‘photopic-dark’, and the second phase ‘scotopic-dark’.

### Suggestive evidence: the dark-background-contingent upshift of gaze

When monkeys fixate targets over dark background, the monkeys appear to fixate a position above the target, not the target itself. For short, we will call this effect ‘the upshift’. Snodderly, who first accidentally discovered it, reported that humans do not show this upshift. Another accidental rediscovery of the upshift was made when the rod-dense region papers were already out. This rediscovery led to a description of some of the upshift’s properties, and to the hypothesis that the effect may have to do with the rod-dense having a role in scotopic vision, the hypothesis under examination in the present study. The next section explicates why the hypothesis was not examined in that study, nor in any other study to date.

A major point of the rediscovery study is that, with a very small target over large uniform background, the upshift appears to be guided by the background, not the target itself. The primary parameter of the background is its absolute luminosity, not the contrast between target and background luminosities. The size of the upshift appears to reflect experience dark-background tasks; a monkey in its first fixation and saccade sessions had small upshift; months later, the same monkey had a larger upshift. At least in some monkeys, the upshift emerges within seconds after the offset of background illumination. Furthermore, we showed that the upshift remains stable through durable fixations, refuting concerns that the upshift reflects the antecedent saccade rather than the fixations themselves.

### Alternative hypothesis: the upshift might be a trait of photopic-dark

The few studies that have addressed the upshift to date have not referred to the state of dark adaptation. Recordings were started briefly after dark onset, because the upshift was thought to be more of an idiosyncrasy than a systematic switching between visual subsystems. Therefore, most of the recordings in studies of the dark-background-contingent upshift were conducted in what we now call photopic-dark. If scotopic-dark had been reached in these studies, it was a rare occssion; certainly full 45-min dark adapatation was rarely if ever actuated. Therefore, even though the hypothesis relates the upshift to scotopic vision, it was not tested if there is an upshift in scotopic vision.

There seem to be two possibilities. Either the hypothesis, tested by the predicition will hold, in which case scotopic vision might have an even larger upshift than the upshift reported with mostly-photopic dark. Or, an alternative hypothesis is that the upshift is not really a feature of scotopic vision. Rather, the upshift emerges in photopic vision; with time in the dark, and the passage to scotopic vision, the upshift would dissipate out out.

### Testing the background requires an ecologically uncommon configuration

In nature, it appears that switching between photopic and scotopic vision occurs almost without exception very slowly, with the time-course determined by that of sunrise and sunset. There are probably hardly ever visual scenes in nature in which a small bright light is superimposed on dark background, certainly dark background after 45-min dark adaptation. Artificial light makes such stimuli available not only in the lab. To set a minimal basis for such stimuli, we will compare them to mesopic lighting.

### Summary

The objective of the present study is to examine the upshift in scotopic conditions, from the viewpoint of an experimental prediction following from the hypothesis the rod-dense superior retina, not the fovea, is used in scotopic vision to fixate the target.

## Methods

Three rhesus monkeys were used in this experiment. All 3 monkeys were adults male, weighting 9-11 kg. All had been previously trained to perform oculomotor tasks for water reinforcement. Hence, the present project required only modest additional training, for accommodating the monkeys to the present task and to dark adaptation.

All experimental procedures are standard and have been described in detail in recent publications of Thier and colleagues (Dash et al., 2012; Caggiano et al., 2013). In brief, eye position was recorded using the scleral search coil method. The monkeys were prepared for neurophysiological recordings (not part of the present study); their heads were painlessly immobilized by titanium head posts. Surgeries were performed under intubation anesthesia with isoflurane and nitrous oxide, supplemented by continuous infusion of remifentanil (1–2.5 (micro)g/kg1·h1) with full tight control of all relevant vital parameters. All procedures conformed to the National Institutes of Health Guide for Care and Use of Laboratory Animals and were approved by the local ethical committee (Regierungspräsidium Tübingen).

The experiments were performed in a standard electrophysiological setup. The monkeys were seated in front of a cathode ray tube (CRT) screen at the distance of 35 cm separating the monkey’s eyes from the center of the screen. The CRT was an Eizo Flexscan F730, 50-cm diagonal, displaying 1,280 × 1,024 pixels at a frame rate of 72 Hz. The room was completely lightproof during the experiment, dark to the level that several human viewers could not report any impression of light after sitting in the closed room for 1 hour. Thus, the only source of light during the experiment was the monitor the monkeys were facing.

Three luminosity levels were used for the background of the target. Bright / photopic background was 7 cd/m2. Dark background (both photopic and scotopic) was 0 cd/m2. For mesopic background we used values between 0.002-0.7 cd/m2. Because we use mesopic vision only to qualitatively compare scotopic background with small bright target, we studied all the mesopic luminosities together.

Two luminosity levels were used for the fixation targets. Small bright targets were 60 cd/m2. Dim targets were 0.01-0.03 cd/m2. In any specific session the dim target luminosity was selected by hand. Initially targets at 0.01 cd/m2 were presented to the monkey; the luminosity was increased until the monkey had started to perform the task reliably. The targets could appear in any of 24 locations, arranged in 3 concentric circles, 8 locations on each circle (Fig 1).

**Figure 1.**
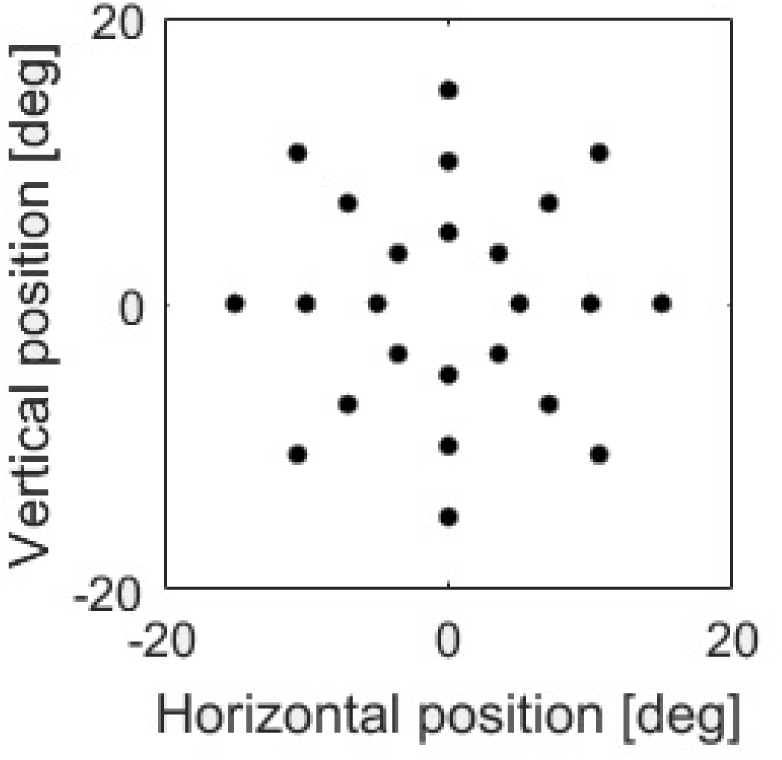
The 24 locations targets could take in this study. The locations are arranged on 3 concentric circles, as illustrated.

The fixation windows used in scotopic vision were 15 degrees radius (horizontal and vertical). In photopic vision the windows were usually 3 degrees, but in some of the trials they were 15 degrees, to verify that the windows do not constrain the monkey’s performance.

The experimental sessions consisted of a series of trial blocks, 120 trials in each block. Interspersed with the blocks were 45-min intervals of dark adaptation (or, some cases, light adaptation, as control, see Results). The mesopic vision condition was tested in a series of blocks that came after the baseline photopic blocks, dark adaptation interval, and scotopic blocks.

Trials lasted for a few seconds (2-5, typically 2-3). The last second of the trial was used for calculating the trial’s median and mean eye position during visual fixation. The search coil data was sampled at a rate of 1000 samples/sec. In a previous study (Spivak et al., 2014) we showed that the mean dark-background-contingent upshift is stable after about 0.5 s from the start of fixation.

Data collection was performed using the open source measurement system nrec, https://nrec.neurologie.uni-tuebingen.de/nrec/nrec, created by F. Bunjes, J. Gukelberger and others, and used as standard in the Thier lab at Tübingen. The nrec output files were converted to Matlab (https://www.mathworks.com/, MathWorks, Natick, Massachusetts, USA), and post hoc data analysis was carried out in Matlab. The statistics used standard t-tests.

## Results

### The results of a session

Data collection spanned multiple daily sessions. By a ‘session’ we refer to the data collected from one monkey, in a single day. A session comprised a series of blocks; a block consisted of a series of trials. In all trials, the monkey’s task was straightforward visual fixation: a very small target (0.02^0^ diameter) appeared on the screen; the monkey made a saccadic eye movement to the vicinity of the target, and remained in the vicinity of the target for as long as the target remained on the screen. At the completion of a predetermined fixation interval (typically, a few seconds) the target disappeared, the monkey was rewarded with a drop of water.

Each trial contained a single target. Target location varied between trials: targets spanned 24 locations (Fig 1), which were presented in a pseudo-random order. Each target appeared several times in a block (typically 4-5). Other than the target’s location, visual parameters, such as target and background luminosities, were maintained throughout each block. Thus, except for an inevitable minimal change of light/dark adaptation occurring throughout a block, the visual conditions were constant throughout each block. Because each block lasted only several minutes, the state of dark adaptation also did not vary significantly in the duration of a block.

Fig 2 shows the results of a typical session including 4 blocks: 2 in photopic and 2 in scotopic conditions. Every block consisted of 120 trials. The two photopic blocks were run first. Then the experimental chamber was totally darkened, and a 45 min dark adaptation period followed, during which no trials took place. After the dark adaptation period was completed, the 2 scotopic conditions blocks were run, one after the other.

**Figure 2.**
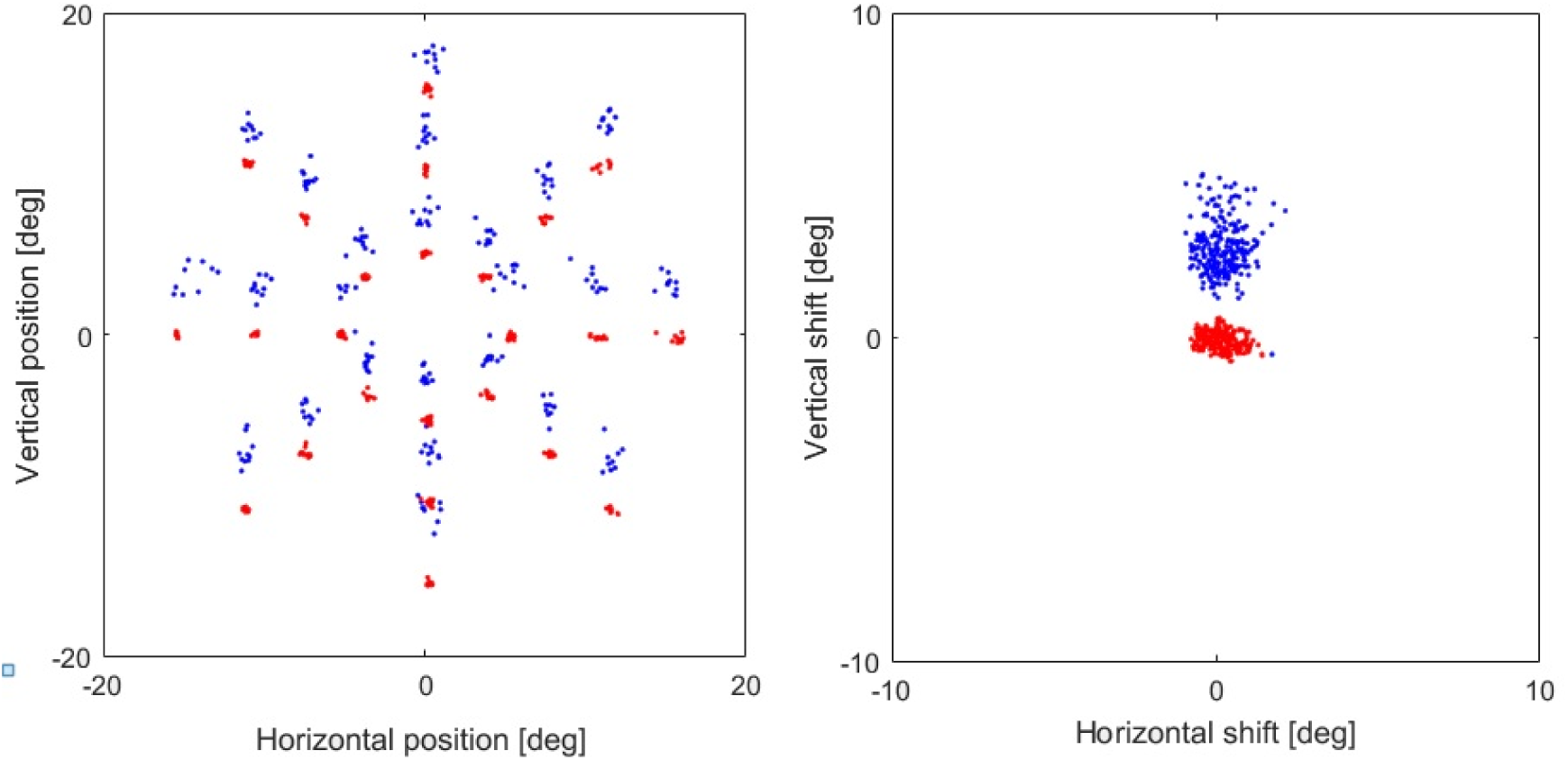
The transformation of trial mean eye position into shift, illustrated for representative blocks. Panel A shows eye position data. Each dot stands for the mean eye position of the fixation in one trial; the dot is plotted in the location the eye had been. Red dots stand for photopic trials (bright target, bright background). Blue dots for photopic-dark background (bright target, dark background). Each red dot cluster is approximately at the position of the target in those trials; each blue dot is above the corresponding red-dot cluster, and is associated with the same target location as the dots in the red cluster. This illustrates the dark-background-contingent upshift previously described. Panel B shows the same data after the transformation to shifts. Red and blue colors retain their significance. Each dot stands for the shift of one trial.

Fig 2A shows the results of the 4 blocks, superimposed. Each dot shows the mean eye position of a single trial. The red dots represent ‘photopic trials’; that is, each red dot depicts the mean fixation position of a trial from the initial two blocks. The blue dots represent ‘scotopic trials’. The red dots are arranged in compact, distinct clusters. The red dots in each cluster reflect trials with the same target position, illustrated in Fig 1: the 3-circle configuration of the targets (Fig 1) is maintained in the configuration of the clusters of the red dots (Fig 2A). The dots in each red cluster are tight, positioned near each other – even though the trials making up a cluster were separated in time by many other trials, directed at targets in other positions, because the 24 targets were presented in a random order. That the overall target configuration (Fig 1) is faithfully reflected by the compact red clusters (Fig 2A), even though the trials of each cluster were scattered over the duration of a block, confirms the precision of photopic fixation.

Importantly, the tightness of the red dot clusters is not caused by external constraints on the deviation of the fixation from target position (‘window size’). See Discussion for a detailed examination of this point. Rather, the tightness of the red clusters reflects the inherent accuracy and precision of photopic fixations.

The blue dots in Fig 2A represent the ‘scotopic trials’; each blue dot depicts the mean fixation position of a trial from the two blocks run after the dark adaptation, with dark background. The blue dots are also arranged in clusters, each cluster corresponding to a target location, similarly, in principle, to the red dot clusters discussed above. Nonetheless, the blue clusters differ from the red clusters, in two important ways. The most prominent difference is in the cluster positions. All blue clusters are visibly shifted to positions above those of the respective red clusters. This is the dark-background-contingent upshift previously described (see Introduction), though here first documented after an appropriate dark-adaptation interval. The second difference is that the blue clusters are visibly less tight than the red clusters. This observation conforms with previous studies of the upshift (see Introduction). Thus, in line with previous studies, we observe in the example session dark-background-contingent upshift, and increase of trial-mean fixation position variability.

### Session analysis: transforming eye position into shift

As Fig 2A illustrates, fixation data were recorded at many target positions over a range in both horizontal and vertical dimensions. This experimental design was primarily aimed to accept results that are not specific to particular target positions. Nonetheless, this experimental design confounds the analysis because eye position cannot be directly used. A look at Fig 2A will help: we want to characterize the relationship between the red dot clusters and the corresponding blue clusters. Evidently eye position cannot be directly used in the analysis, because it varies from one red cluster to the next. To resolve this confound, we define ‘shifts’, which are invariant of target positions. For the trials of the example session, Fig 2 illustrates the transformation of the mean eye-position of all the session’s trials (Fig 2A) into shifts (Fig 2B). Each red dot in Fig 2A corresponds to a red dot in Fig 2B, and each blue dot in Fig 2A corresponds to a blue dot in Fig 2B.

#### Definition of shift

The rationale guiding the definition of shift is the following. The shift is to be the outcome of a calculation whose input is eye-position records. For the shift outcome to be independent of target position, the calculation itself must depend on target position. The calculation must first compare separately for each target position the scotopic and photopic eye position, or the blue dot cluster and the red dot cluster, both corresponding to the target at stake; and then generalize over all target positions.

Towards this aim we define, for each target position, a reference photopic fixation position. We define it as the mean of the red dot cluster at stake. Keeping in mind that each red dot represents a mean fixation position, of all the eye-position samples in the time-interval of the relevant trial, it transpires that the reference position is a mean of means. Each trial’s mean position contributes equally to the reference position, with the horizontal and vertical dimensions calculated separately.

Given a fixation target, and the reference eye-position associated with that target, we now define the shift. The shift is the vectorial difference of the eye-position at stake and the reference eye position. This definition allows to transform both isolated eye-position samples and the mean eye-position of given trials into shifts. As long as the target is well defined, so is the transformation from eye-position samples to shifts.

With this definition, the disparate mean eye position red and blue clusters of Fig 2A are transformed to a small cluster of red shifts all close to (0,0) and a larger cluster of blue shifts above the red cluster, illustrated in Fig 2B. Thus the shifts appear to allow comparing data from all trials together, regardless of target position.

We define ‘upshift’ as the vertical components of the shift. This term reflects previous findings that scotopic fixation positions tend to be above photopic. Thus, upshift is defined for all entities for which shift is defined. Here we will refer mainly to mean upshifts of trials.

#### Verification that the shift reflects all target positions, not outliers

Fig 3 illustrates the distributions of the vertical components of the mean eye position of the trials of the example session – the vertical dimension of the dots in Fig 2A – but with respect to their reference fixation positions. The photopic trials are centered very close to zero, in line with the definition of the reference position; the mean and standard deviation are 0.08^0^±0.2^0^ degrees. The scotopic trials’ mean positions are shifted upwards and are more variable, at 2.72^0^±0.8^0^ degrees. The two distributions are almost entirely disjoint (there is one overlapping bin drawn in white, close to 0). Hence, by using each target’s reference position, a global difference between all scotopic trials and all photopic trials is revealed (Fig 3), invariant of each trial’s target position.

**Fig 3.**
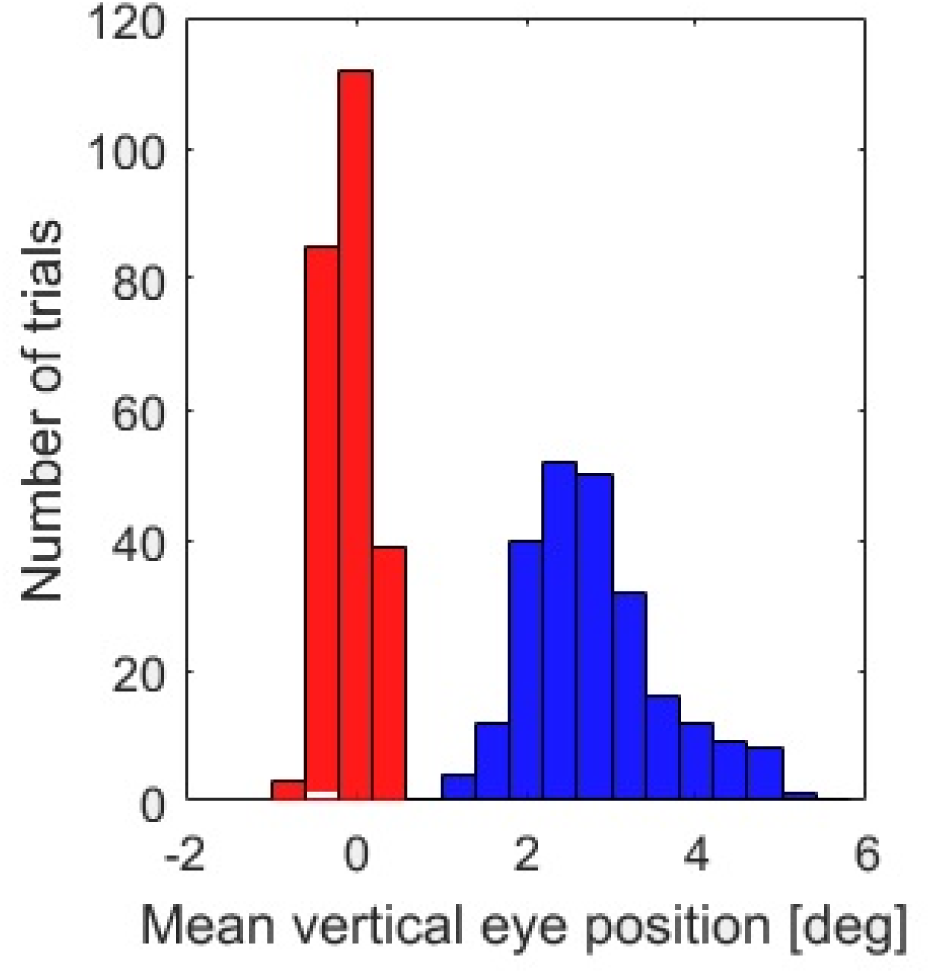
Histograms of the vertical components of the mean eye position of the trials of the example session – the vertical dimension of the dots in Fig 2A – drawn with respect to their reference fixation positions. See the main text for definition of reference eye positions.

By using the reference position as anchor, the mean upshift can be computed for each target position. The distribution of the upshifts at each of the 24 target positions is illustrated for the example session

The justification for this lumping together comes from the observation that in each of the target locations the red and blue dots are entirely separated. Indeed, the means significantly differ between the vertical components of the blue and red dots: the mean vertical eye position in scotopic conditions was 2.720±0.80 and in photopic conditions: −0.080±0.20. In all target positions the difference of the blue and red means is significant (p values ranging between 9e-14 to 1.2e-5). The difference between scotopic and photopic horizontal components was mainly not significant except for the two most eccentric horizontal targets (p = 0.037 and p = 0.048 for the right and for the left target respectively). Further, Fig 3 shows a histogram of the means of the vertical positions and superimposed the mean of the entire session, depicted in Fig 2B. Fig 4 shows the distribution of upshift over 24 possible target locations. Clearly the mean is not determined by extreme points.

**Fig 4.**
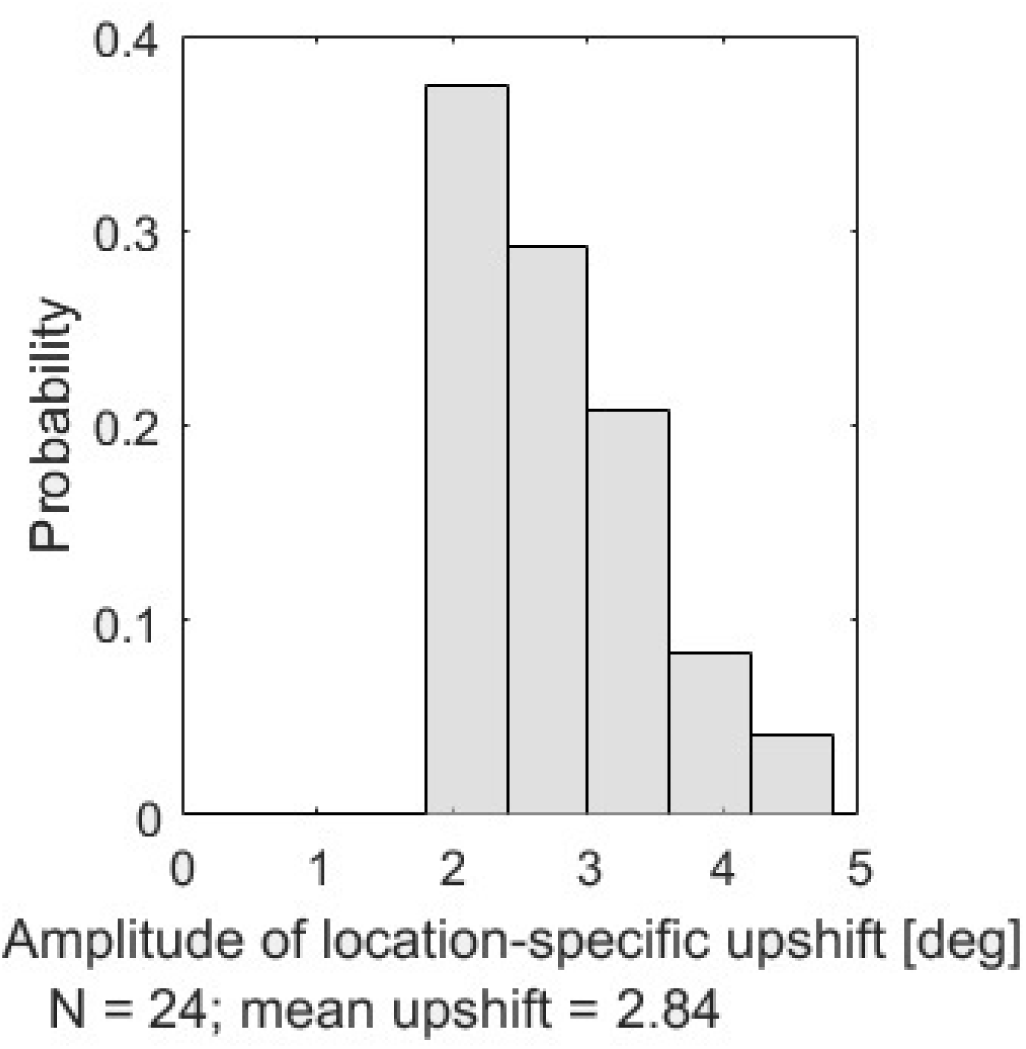
Histogram of mean upshifts, calculated separately for the 24 target locations, shows that the transformation into shifts is representative of the entire field.

### Modular analysis

In the following we will repeatedly compare upshifts in pairs of conditions, and will use the same format to present the results. The present section describes the format, using Fig 5 as reference. Each Monkey’s results are depicted in a row of panels; Fig 5A,B illustrate Monkey P’s results, Fig 5C,D Monkey L’s. In some cases there are results from 3 monkeys, organized in the rows of panels. The results of each monkey are presented as scatterplot on the left (Figs 5A,C), and histograms on the right (Fig 5B,D). The scatterplots show 2-dimensional shifts. Each dot stands for a single trial; the dot is placed at the coordinates corresponding the mean shift of that trials. Dots are in one of 2 or 3 colors. Red dots tend to stand for trials in photopic conditions (bright target, bright background); blue for dark (dark background, various variations). The same number of dots are displayed for each color; the selection of trials to be displayed is random for the more populous condition. Nonetheless, because of the large number of trials the present study involves, dots often overlap each other, occluding the shapes of the distributions. Thus, presented on the right are histograms of the upshifts, the vertical components of the shifts. The histograms are plotted with respect to the vertical axis of the scatterplot. The colors of the dots are conserved in the histograms; overlapping bins are presented in white.

**Fig 5.**
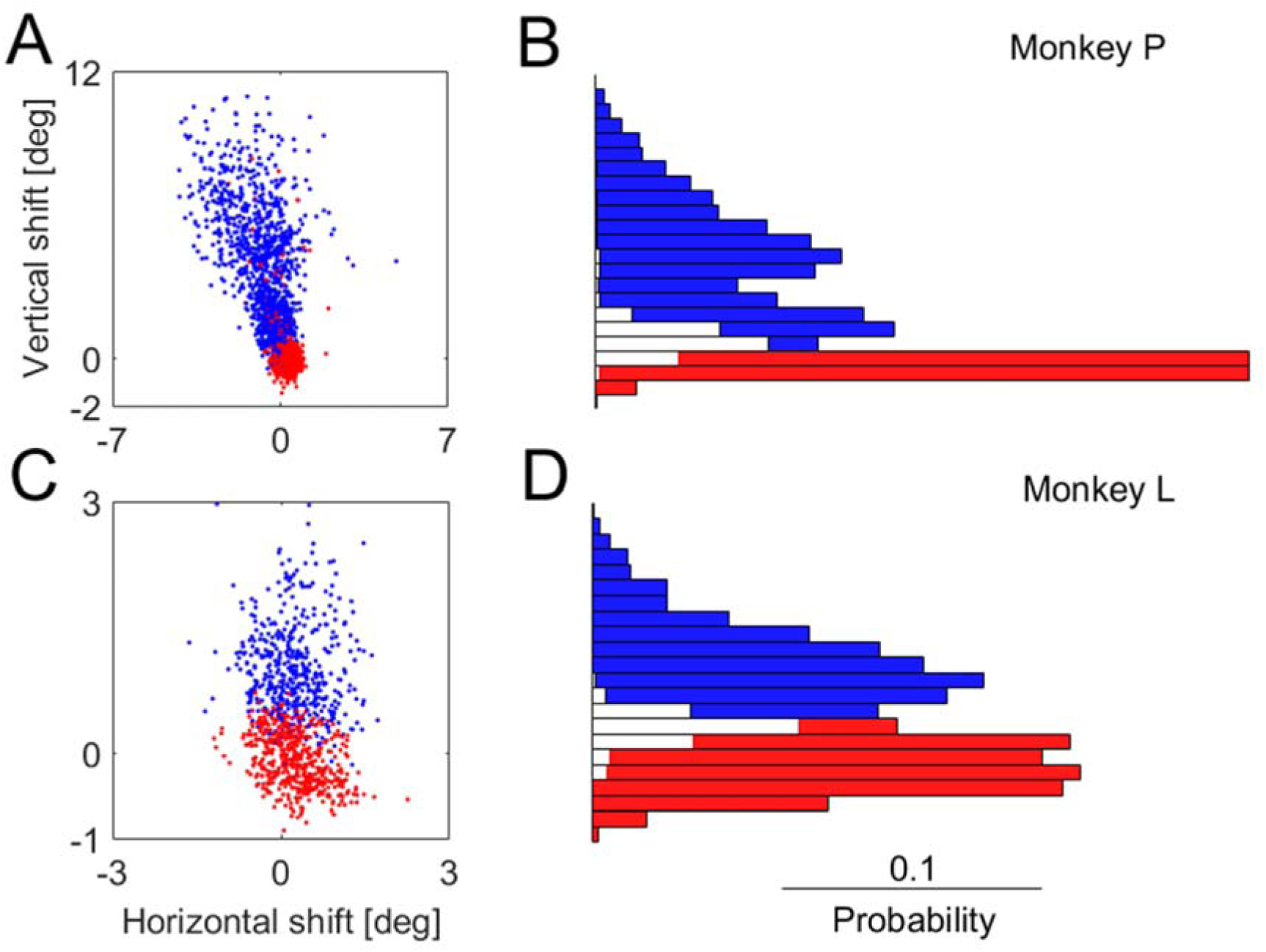
Within-session analysis of the upshift in photopic-dark background (blue) compared to photopic (red). Each row of panels illustrates the data of one monkey. The monkey’s code is listed on the right side of the row. On the left (panels A,C) are scatterplots of the shifts of the monkey’s data. Each dot stand for one trial, with the color coding the condition of the trial (see above). The vertical dimension of the dots in the scatterplot (and possibly more data, see main text) are analyzed in the histograms on the right (panels B,D). White bins in the histogram stand for overlap of the blue and red. Table 1 contains the relevant statistics.

**Table 1.**
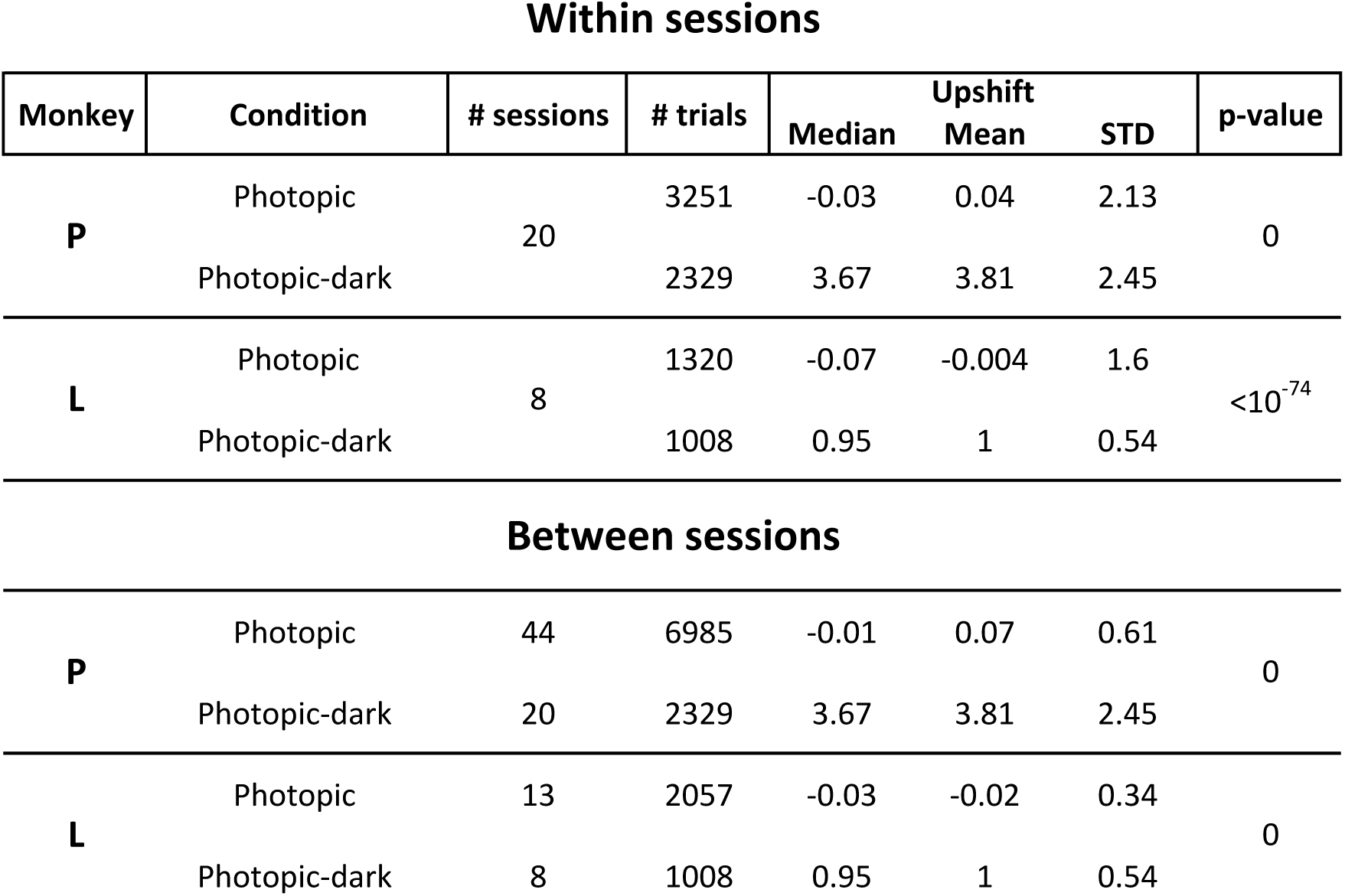
Photopic-dark background versus photopic vision

Within sessions
| Monkey | Condition | # sessions | # trials | Median | Upshift Mean | STD | p-value |
| --- | --- | --- | --- | --- | --- | --- | --- |
| **P** | Photopic | 20 | 3251 | -0.03 | 0.04 | 2.13 | 0 |
|  | Photopic-dark |  | 2329 | 3.67 | 3.81 | 2.45 |  |
| **L** | Photopic | 8 | 1320 | -0.07 | -0.004 | 1.6 | <10<sup>-74</sup> |
|  | Photopic-dark |  | 1008 | 0.95 | 1 | 0.54 |  |
| Between sessions |  |  |  |  |  |  |  |
| **P** | Photopic | 44 | 6985 | -0.01 | 0.07 | 0.61 | 0 |
|  | Photopic-dark | 20 | 2329 | 3.67 | 3.81 | 2.45 |  |
| **L** | Photopic | 13 | 2057 | -0.03 | -0.02 | 0.34 | 0 |
|  | Photopic-dark | 8 | 1008 | 0.95 | 1 | 0.54 |  |

The statistics of each comparison are presented in an associated table; for Fig 5 it is the top part of Table 1. For each monkey, separately, the numbers of sessions and trials are presented, as well as the mean, standard deviation, and median of the upshift, in degrees of visual angle. Finally, the p-value of the t-test comparing the means of the two distribution is listed. Zero (0) indicates that the p-value is less than the smallest value representable by standard computer double-precision arithmetic.

### Within-sessions versus between-sessions comparisons

Before indulging onto the body of the results we need to make a methodological distinction. Ideally, we can compare the upshifts in two conditions run one after the other, within the same session. For example, in previous studies describing the upshift a first condition was fixation of bright targets over bright background (photopic vision). Then, shortly after dark onset, fixation in a second condition was collected (bright targets over dark background, shortly after dark onset). That these conditions could be collected within the same session meant that they shared the same eye-positon calibration.

When 45-min waiting periods have to be had, monkeys are generally willing to work after the period ends. (This holds regardless of whether the room is lighted or dark during the waiting period). However, a second waiting period is just too much. Because each condition typically involves collection of hundreds of trials, in general during a session we could record two conditions, one, baseline, before a waiting period, usually 45-min dark adaptation; the other after the waiting period.

The need to make one comparison, with one waiting period, per session, constrained the comparisons we could make within sessions. Consequently, in some cases we had to make between-session comparisons. Transforming eye-position data into shifts makes that possible. Since different sessions have different calibrations, we assume that between-session comparisons might be more noisy. However, as described subsequently, all the between-sessions comparisons we make are highly significant in all monkeys tested.

We always have more between-session data than within-session. Therefore, in the comparisons for which we have within-session data, we describe these comparisons. In all cases we describe also the between-session comparisons with all the available data.

### Photopic-dark background evokes upshift

**At issue:** Does photopic-dark background evoke upshift? Previous studies, which documented the dark-background-contingent upshift, probably tested eye position mostly in photopic-dark, but this was not well controlled as the state of dark adaptation was not documented.

**Design:** We compare two conditions. Condition 1 comprises ‘photopic’ trials. The target is very small (0.02^0^), only a few pixels on the computer monitor, but bright enough to be salient over the bright background (see Methods of parameters). Condition 2 comprises ‘photopic-dark background’ trials. The same target, size and luminance as in condition 1, but the background is dark. The entire photopic-dark data is collected during the photopic leg of the dark adaptation curve, during the first 10 min after dark onset.

The null hypothesis states that the distributions of upshifts in photopic and in photopic-dark background conditions would be the same; hence, their means would not differ significantly. With the target kept unchanged between the conditions, any difference in upshift would reflect the changing background, not the target.

**Results:** Data were collected from 2 monkeys. Fig 5 illustrates the within-session results; Fig 6, the between-sessions. Table 1 specifies the statistics. The Figures follow the pattern described in the previous sections.

**Fig 6.**
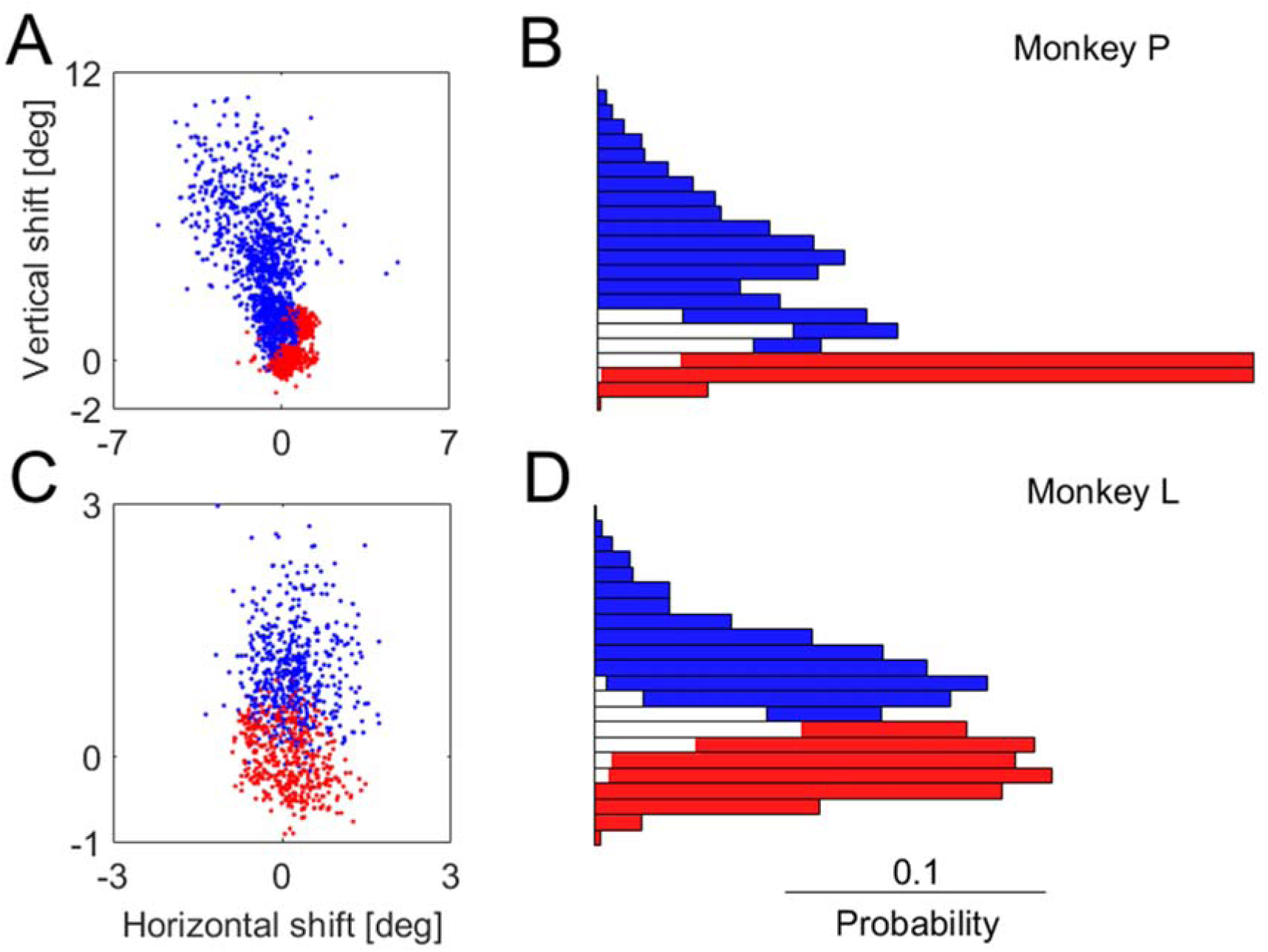
Between-session analysis of the same categories as Fig 5 showed for within-data analysis. Table 1 contains the statistics. Note the much larger number of trials involved in the present analysis, but the higher variability typical of between-session analysis.

In both monkeys, the photopic-dark (blue) and photopic (red) trials make up two largely distinct clusters of dots in the scatterplots, and distributions in the histograms. Monkey P has a greater upshift than Monkey L, but in Monkey L, as well as in Monkey P, the means of the upshifts in the red and blue trials are significantly different, with a *p-value* of zero (less than the smallest value representable by standard computer arithmetic).

**Conclusion:** The null hypothesis is rejected. Thus, fixation positions in photopic-dark are shifted above those in photopic vision.

### Scotopic-dark background evokes upshift

**At issue:** In the previous section, we observed that photopic-dark background evokes upshift. Does scotopic-dark background at all evokes upshift, too? The working hypothesis of this study contends that it will; an alternate hypothesis could state that the upshift phenomenon is limited in time to the photopic leg (first 10-15 min) of darkness; subsequently, the upshift would dissipate. Because previous studies did not explore the upshift in scotopic-dark, the issue is open.

**Design:** The compared conditions are almost identical to those in the previous section. Condition 1 is photopic, bright target over bright background; condition 2 is bright target over dark-background, as in condition 1 – but with one difference: after the monkey completes the photopic condition blocks, the experimental chamber is completely darkened for a 45-min interval of dark adaptation. Only then the monkey is presented with, and performs, the dark-background blocks. We call this condition ‘scotopic-dark background’. Inevitably, the use of a bright target, however small, compromises the state of dark adaptation. Nonetheless, because the target is very small, this compromising effect is not very large. The advantage of using this combination, of a very small bright target over scotopic-dark background, is that we can compare directly the effect of the dark adaptation specifically of the background, in isolation from the target. This is explored in the next section.

Thus, our working hypothesis for this section is that there is upshift in the scotopic-dark background conditions. The null hypothesis states that the distributions of upshifts in the two conditions are the same, hence their means are also the same.

**Results:** Data were collected from 2 monkeys. Fig 7 illustrates the within-session results; Fig 8, the between-sessions. Table 2 specifies the statistics. The Figures follow the pattern described in the previous sections. In both Figures, red stands for photopic trials (bright target over bright background); blue stands for scotopic-dark background (bright targets over dark background, after 45-min dark-adaptation interval).

**Fig 7.**
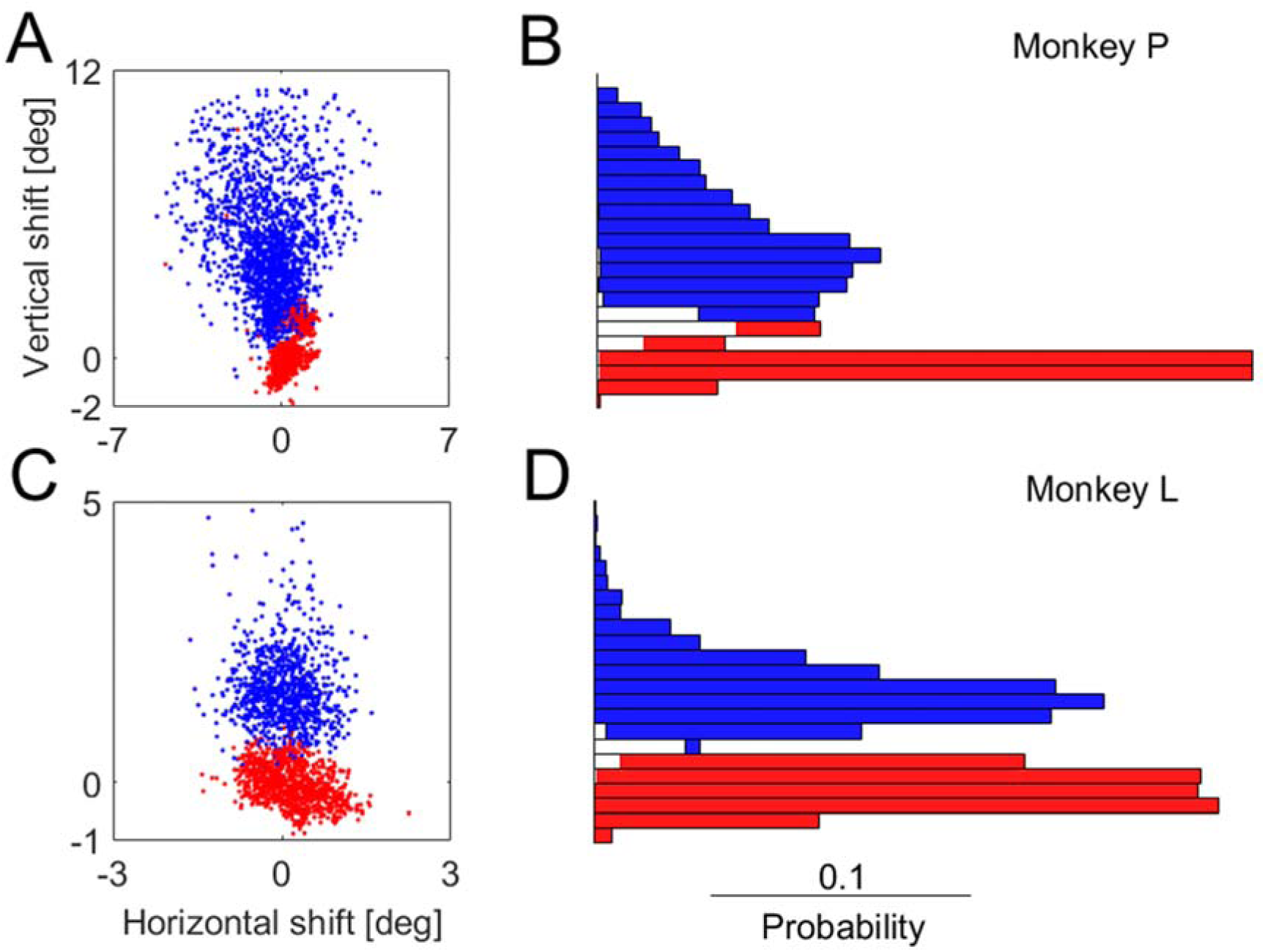
Within-session analysis of the upshift in scotopic-dark background (blue) compared to photopic (red). Same format as Fig 5. Statistics included in Table 2.

**Fig 8.**
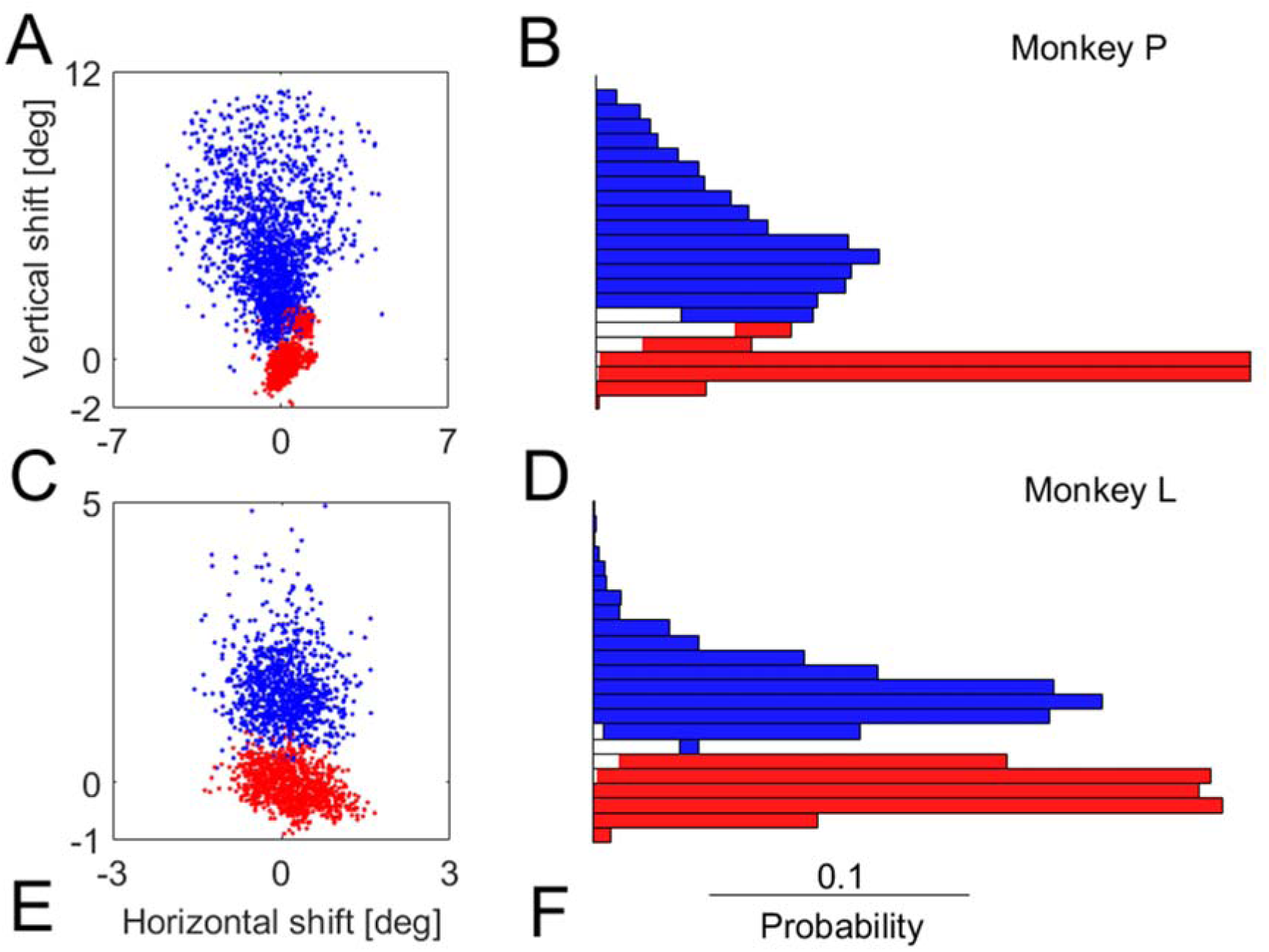
Between-session analysis of the same comparison as Fig 7. Same format as Fig 5. Statistics included in Table 2.

**Table 2.**
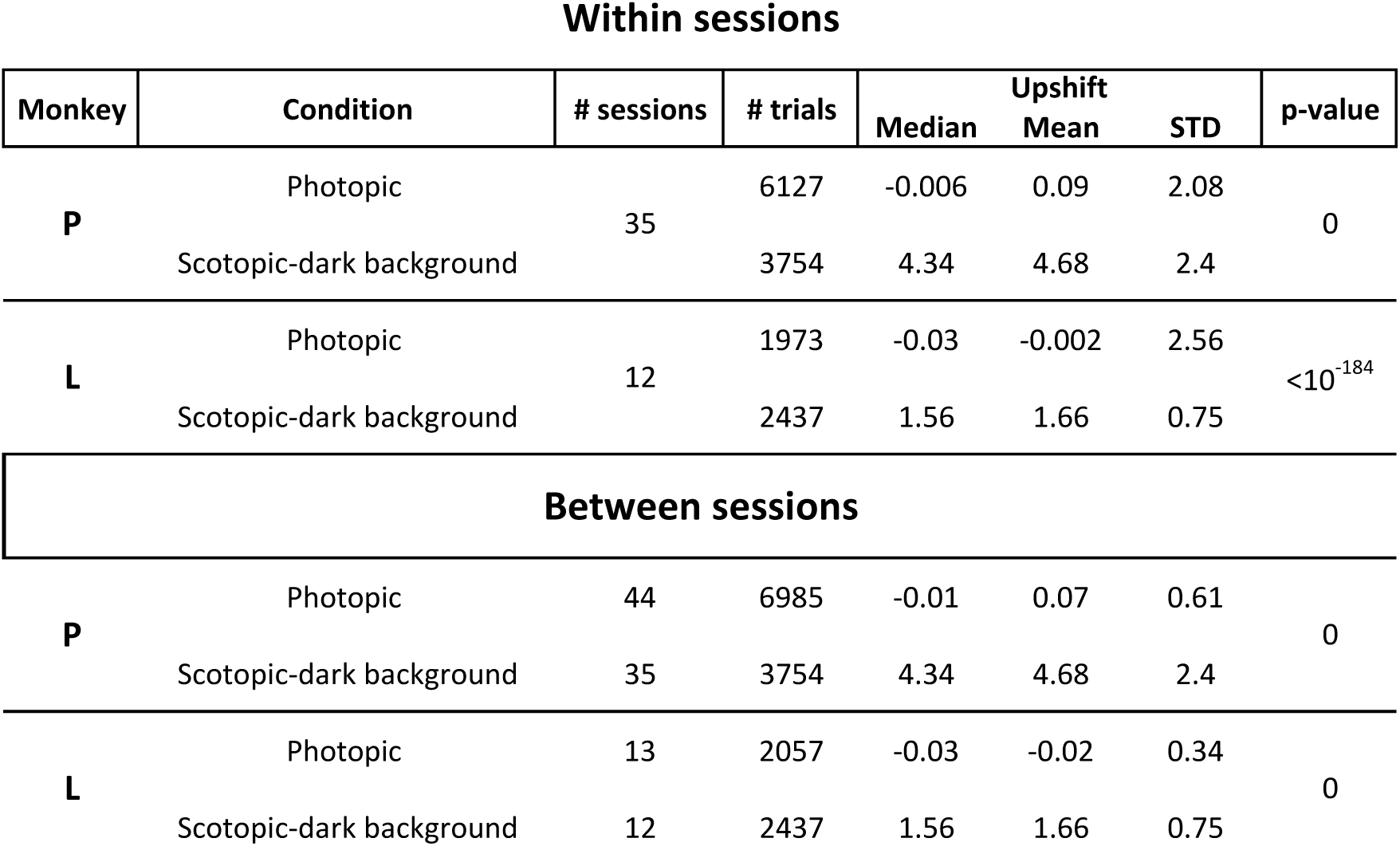
Scotopic-dark background versus photopic vision

In both monkeys, blue and red dots in the scatterplots are visibly almost totally separate, with the blue dots clusters centered almost precisely above the red dot clusters. The blue clusters of Monkey P are elongated in the vertical direction, in both Figs 7 and 8; this is reflected in the higher standard deviation (see Table 2). The mean and median upshift values are greater in monkey P than in Monkey L. Nonetheless, in both monkeys, the means are significantly different, with a *p-value* of 0.

**Conclusion:** The null hypothesis is rejected. There is dark-background-contingent upshift after the 45-minute dark adaptation interval, which converts the dark to scotopic. Thus, the upshift is not specific to photopic-dark.

### A greater upshift is evoked by scotopic-dark background than by photopic-dark background

**At issue:** We now know that there is upshift with both photopic-dark and scotopic-dark backgrounds. Is the upshift equal in both conditions? If not, how do the two relate? At the background is the question whether the upshift is associated primarily with one of these phases in the process of dark adaptation, photopic or scotopic. A larger upshift in one of the two would suggest a special relationship of that stage to the upshift.

**Design:** The shifts of the photopic-dark trials cannot be directly compared with the shifts of the scotopic-dark trials. There are no sessions containing both photopic-dark background and scotopic-dark background trials. (Had we done that, we would have broken the standard structure of the sessions, possibly confounding the results in other ways). Therefore, the comparison is possible only in the between-sessions mode. Accordingly, the working hypothesis is that the upshift is greater in the scotopic-dark background than in the photopic-dark background trials.

The null hypothesis states that the two states of darkness lead to the same distribution of upshifts; or, more specifically, that photopic-dark evokes the same or greater upshift than scotopic-dark.

**Results:** The data compared here overlaps that presented in Figs 5-8. Hence, the data are from the same 2 monkeys. Fig 9 illustrates the between-sessions comparison. Table 3 specifies the statistics. The Figure follows the pattern described in the previous sections. Green stands for photopic-background trials (bright target over dark background, without dark adaptation); blue stands for scotopic-dark background (bright targets over dark background, after 45-min dark-adaptation interval). Recall that white in the histogram stands for overlap of the two distributions.

**Fig 9.**
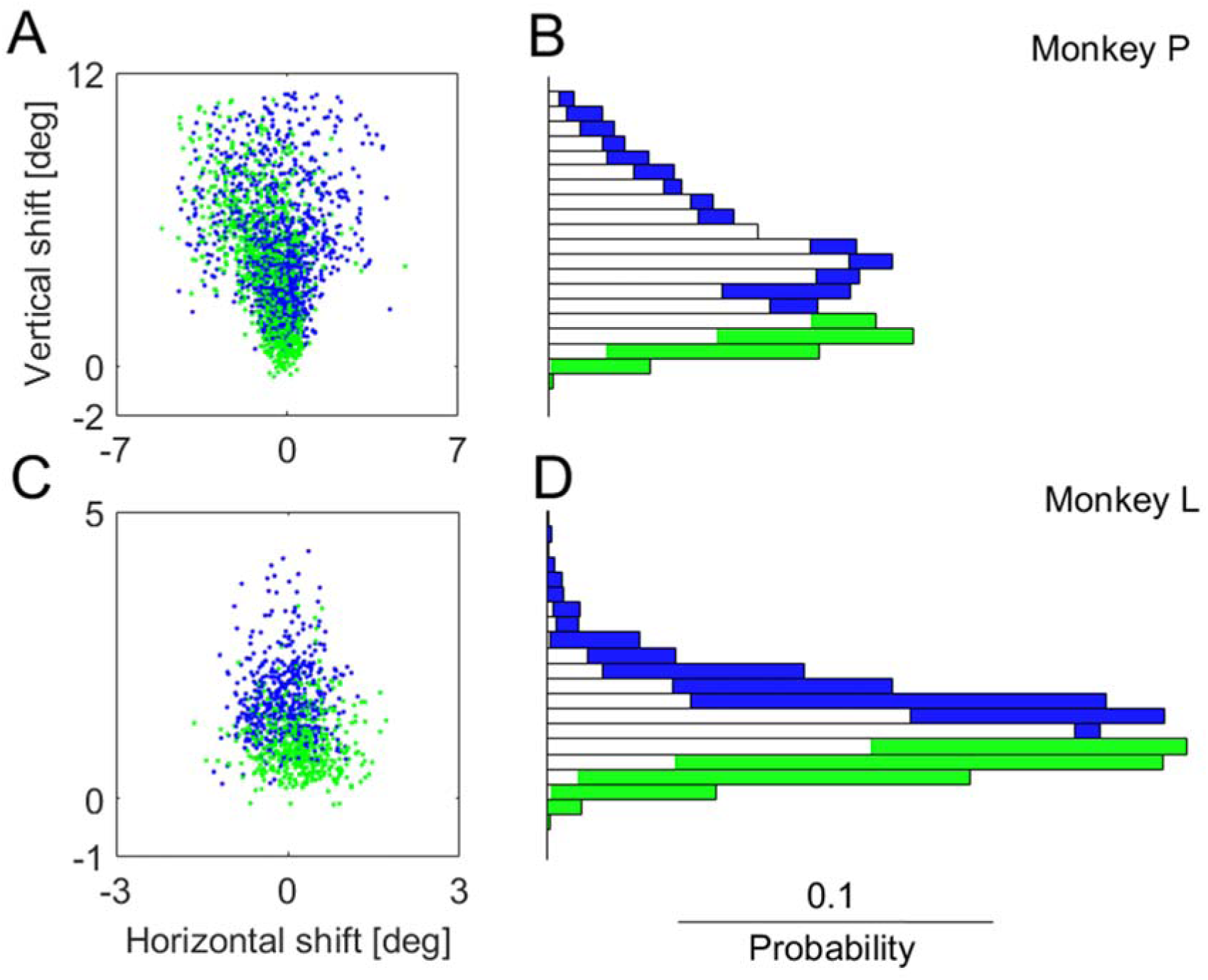
Between-session analysis of the upshift evoked by scotopic-dark background, compared to photopic-dark background. Same format as Fig 5. Statistics included in Table 3.

**Table 3.** Scotopic-dark background versus photopic-dark background

Between sessions
| Monkey | Condition | # sessions | # trials | Median | Upshift<br>Mean | STD | p-value |
| --- | --- | --- | --- | --- | --- | --- | --- |
| **P** | Scotopic-dark background | 35 | 3754 | 4.34 | 4.68 | 2.39 | $>10^{-41}$ |
|  | Photopic-dark | 20 | 2329 | 3.67 | 3.81 | 2.45 |  |
| **L** | Scotopic-dark background | 12 | 2437 | 1.56 | 1.66 | 0.75 | $>10^{-126}$ |
|  | Photopic-dark | 8 | 1008 | 0.95 | 1 | 0.53 |  |

In both monkeys, there is large overlap between the green and blue data. This is easily visible in both scatterplots (Figs 9A,C) and histograms (figs 9B,D). The overlap between the blue and green histograms is much larger than the overlap between the red and blue histograms in Figs 5-8.

Nonetheless, especially in the histograms but also in the scatterplots, it is apparent that the blue dots and bins tend to lie above the green dots and histogram bins. That holds for both monkeys. The means of the distributions are very significantly different, with *p-value* less than 10^-41^ in both cases. Because the *p-value* is so small, both variations of the null hypothesis are rejected.

**Conclusion:** The null hypotheses are rejected. In scotopic-dark background, a larger upshift is evoked than in photopic-background.

### Scotopic vision evokes upshift

**At issue:** Is there upshift in scotopic vision? We know now that bright targets over scotopic-dark background evoke upshift; could the bright targets be necessary for evoking the upshift? The combination of a bright target over scotopic-dark background is an ecologically rare stimulus. Is the upshift but an idiosyncratic response to this ecological rarity?

**Design:** Condition 1 is photopic vision, bright target over bright background. Condition 2 is scotopic vision, dim target over dark background, after 45-min dark adaptation. Both the scotopic-background condition, described in the previous 2 sections, and the current full scotopic-vision condition, involved the same 45-min dark-adaptation interval, and the same dark background, but differed in the target luminosity. In the dark-background condition, the target was bright; in the current, scotopic-vision condition, the target was dim, 0.01 cd/m^2^ or a little more, if required for the monkey to perform his task.

The null hypothesis is that scotopic vision does not evoke upshift; thus, both photopic and scotopic vision conditions would involve the same distribution of upshifts, with the same means.

**Results:** Data were collected from 3 monkeys. Fig 10 illustrates the within-session results; Fig 11, the between-sessions. Table 4 specifies the statistics. The Figures follow the pattern described in the previous sections. In both Figures, red stands for photopic trials (bright target over bright background); blue stands for scotopic (dim targets over dark background, after 45-min dark-adaptation interval).

**Fig 10.**
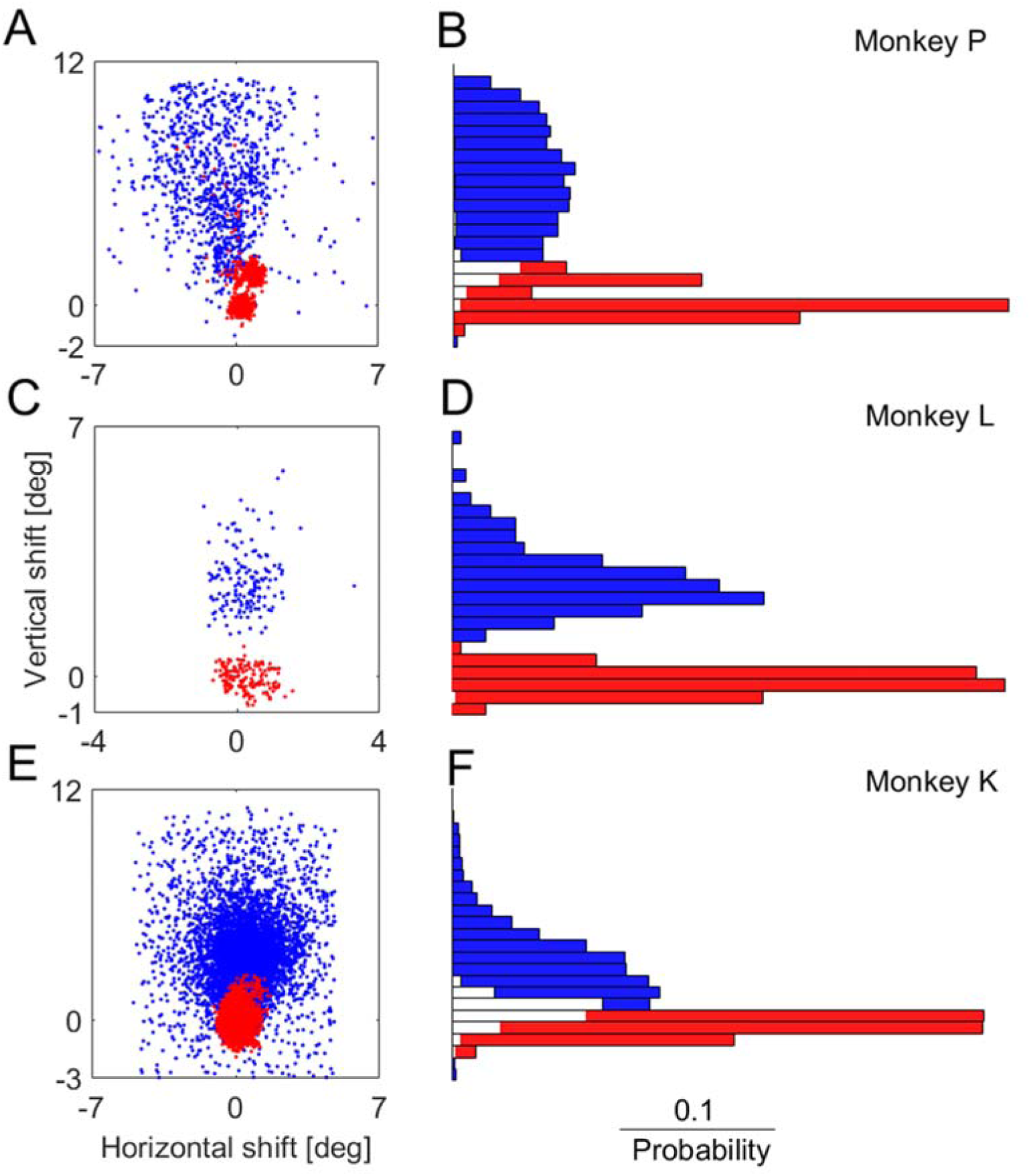
Within-session analysis of the upshift in scotopic vision (blue) compared to photopic (red). Same format as Fig 5. Statistics included in Table 4.

**Fig 11.**
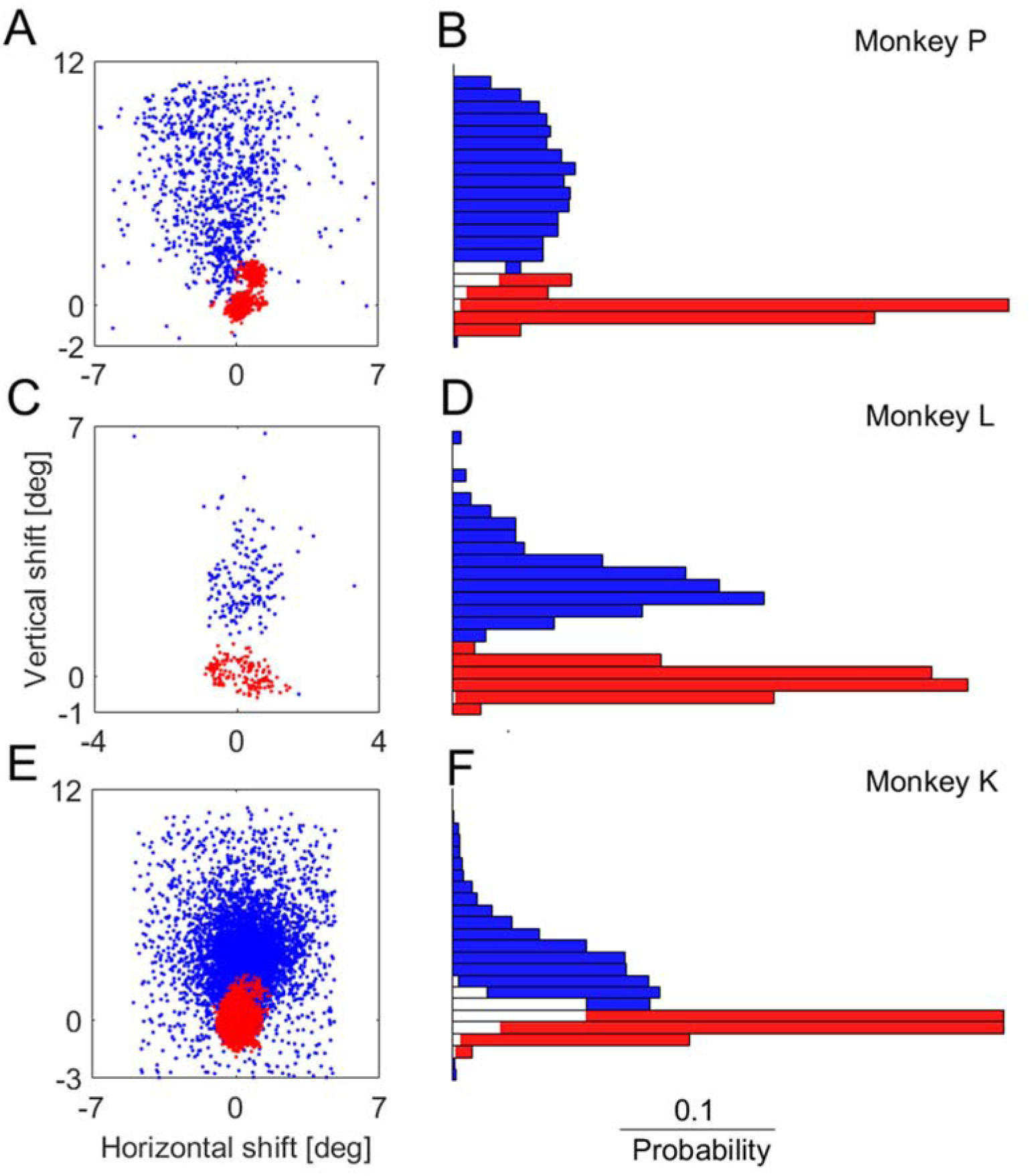
Between-session analysis of the same comparison as Fig 10. Same format as Fig 5. Statistics included in Table 4.

**Table 4.**
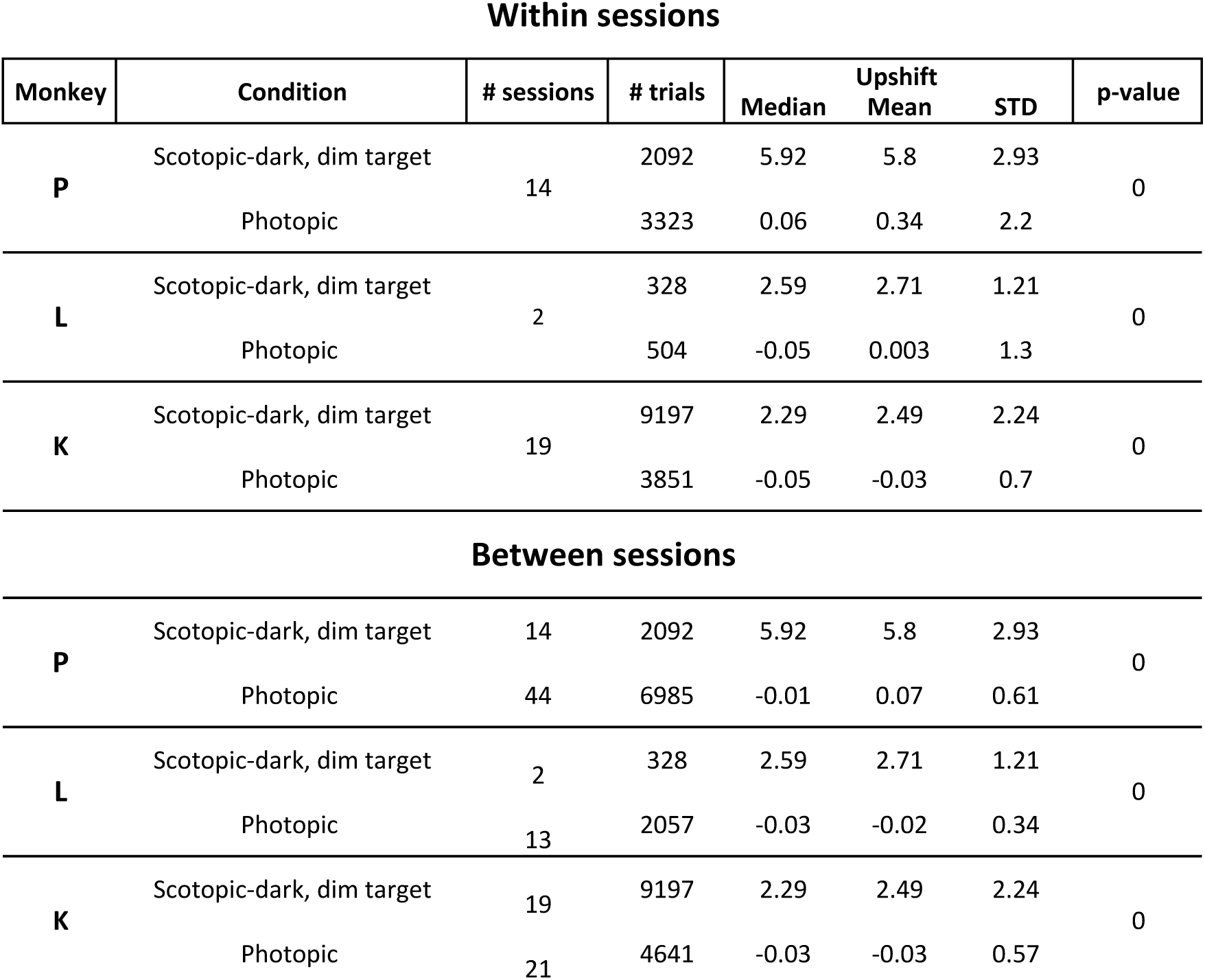
Scotopic vision versus photopic vision

Within sessions
| Monkey | Condition | # sessions | # trials | Median | Upshift Mean | STD | p-value |
| --- | --- | --- | --- | --- | --- | --- | --- |
| **P** | Scotopic-dark, dim target | 14 | 2092 | 5.92 | 5.8 | 2.93 | 0 |
|  | Photopic |  | 3323 | 0.06 | 0.34 | 2.2 |  |
| **L** | Scotopic-dark, dim target | 2 | 328 | 2.59 | 2.71 | 1.21 | 0 |
|  | Photopic |  | 504 | -0.05 | 0.003 | 1.3 |  |
| **K** | Scotopic-dark, dim target | 19 | 9197 | 2.29 | 2.49 | 2.24 | 0 |
|  | Photopic |  | 3851 | -0.05 | -0.03 | 0.7 |  |
| Between sessions |  |  |  |  |  |  |  |
| **P** | Scotopic-dark, dim target | 14 | 2092 | 5.92 | 5.8 | 2.93 | 0 |
|  | Photopic | 44 | 6985 | -0.01 | 0.07 | 0.61 |  |
| **L** | Scotopic-dark, dim target | 2 | 328 | 2.59 | 2.71 | 1.21 | 0 |
|  | Photopic | 13 | 2057 | -0.03 | -0.02 | 0.34 |  |
| **K** | Scotopic-dark, dim target | 19 | 9197 | 2.29 | 2.49 | 2.24 | 0 |
|  | Photopic | 21 | 4641 | -0.03 | -0.03 | 0.57 |  |

In all 3 monkeys, in both within-sessions and between-sessions analysis, the scotopic vision fixations lie above the photopic fixation positions, with very small overlap. The values of the scotopic upshifts vary considerably, within monkey and between monkeys; in all monkeys the standard deviation of the scotopic upshifts is 3.5-5 times the standard deviation of the photopic upshifts. Thus, almost all scotopic fixations are above the range of photopic fixations. The size of the shift varies between trials and between monkeys. The trial-by-trial variability of the size of the upshift is a major aspect of the results, and will be resorted to in the Discussion.

As for the null hypothesis: the means of the scotopic and photopic distributions are significantly different, with *p-value* 0.

**Conclusion:** The null hypothesis is rejected. In scotopic vision, there is upshift.

### A greater upshift is evoked by full scotopic vision than by scotopic-dark background

**At issue:** We know now the upshift is present in scotopic-dark background, with a bright small target, and in scotopic vision proper, with scotopic-dark background and dim target. Thus, the alternate hypotheses that related the upshift to other factors than scotopic vision – photopic-dark, the ecological rarity of a bright target over scotopic background – failed to explain the data. The central hypothesis of this study, that the upshift is associated with scotopic vision because it reflects a tendency to fixate by the rod-dense region in superior retina, is still intact.

To give this hypothesis greater strength, the prediction that full scotopic vision involves a greater upshift than scotopic-dark background should hold.

**Design:** This comparison is possible only between sessions, because the two conditions follow 45-min dark adaptation intervals. Condition 1 is scotopic-dark background (bright target, dark background, after 45-min dark adaptation). Condition 2 is full scotopic vision (dim target, dark background, after 45-min dark adaptation). The null hypothesis states that the distributions of upshifts in the two conditions are the same; hence, their means are equal.

**Results:** Fig 12 shows the results, and Table 5 details the statistics. The format is the same as in previous Figures. Green stands for scotopic-dark background, blue for full scotopic vision. White histogram bins represent the overlap between the blue and green histograms.

**Fig 12.**
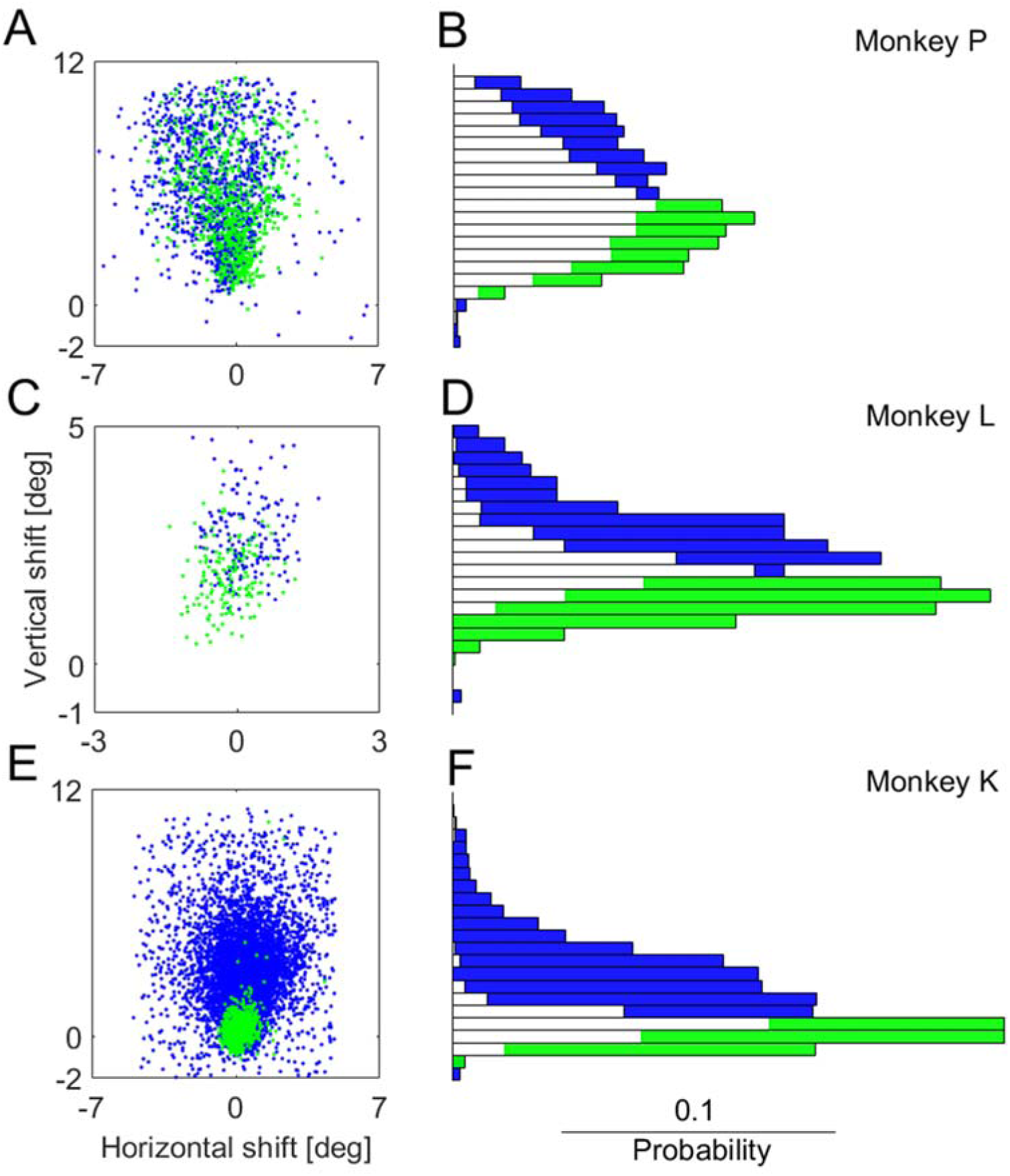
Between-session analysis of the upshift evoked by scotopic-dark background, compared to full scotopic vision.

**Table 5.** Full photopic vison versus scotopic-dark background

Between sessions
| Monkey | Condition | # sessions | # trials | Median | Upshift Mean | STD | p-value |
| --- | --- | --- | --- | --- | --- | --- | --- |
| **P** | Scotopic-dark, dim target | 14 | 2092 | 5.92 | 5.8 | 2.93 | $>10^{-54}$ |
|  | Scotopic-dark background | 35 | 3754 | 4.34 | 4.68 | 2.39 |  |
| **L** | Scotopic-dark, dim target | 2 | 328 | 2.59 | 2.71 | 1.21 | $>10^{-95}$ |
|  | Scotopic-dark background | 12 | 2437 | 1.56 | 1.66 | 0.75 |  |
| **K** | Scotopic-dark, dim target | 19 | 9197 | 2.29 | 2.49 | 2.24 | $>10^{-201}$ |
|  | Scotopic-dark background | 2 | 1078 | 0.3 | 0.35 | 0.69 |  |

The amount of overlap varies between monkeys. It is greatest in Monkey P, intermediate in Monkey L, and small in Monkey K. Interestingly, the monkey with the largest upshift (P) has the greatest overlap. We will refer to this observation in the Discussion.

In all 3 monkeys, the blue dots and histogram bins are evidently overall positioned above the green ones. The means are very significantly not equal, with *p-value* of 10^-54^ or less in all monkeys (see Table 5).

**Conclusion:** Scotopic vision evokes upshifts that are higher than those evoked by scotopic-dark background, consistent with the central tenet of this study.

### Mesopic vision evokes upshift

**At issue:** An important aspect of the results, presented in the previous sections, is that in many conditions there is an upshift, but its size is not maximal. Bearing in mind that some of these conditions comprise ecologically questionable visual stimuli (the combination of a bright target and completely dark background), we ask if an ecologically common stimulus can also evoke upshift, but whose level is less than maximal. Mesopic-background stimuli are called for. Common in nature, intermediate between scotopic and photopic, we could expect the upshift to be sub-maximal. In the present section we test if there is upshift in the mesopic blocks.

**Design:** Condition 1 is photopic vision, bright target over bright background, as used before. After the photopic vision blocks, the monkeys went through a 45-min dark adaptation interval. After some testing in scotopic-background conditions, the background was changed to a mesopic level (see Methods for details). A standard small, bright target and mesopic background were used in the ‘mesopic’ blocks of trials that followed.

The null hypothesis predicts there is no change in upshift between photopic and mesopic blocks.

**Results:** Data were collected from 2 monkeys. Fig 13 illustrates the within-session results; Fig 14, the between-sessions. Table 6 specifies the statistics. The Figures follow the pattern described in the previous sections. In both Figures, red stands for photopic trials (bright target over bright background); cyan stands for mesopic (bright target over mesopic background, as described above).

**Fig 13.**
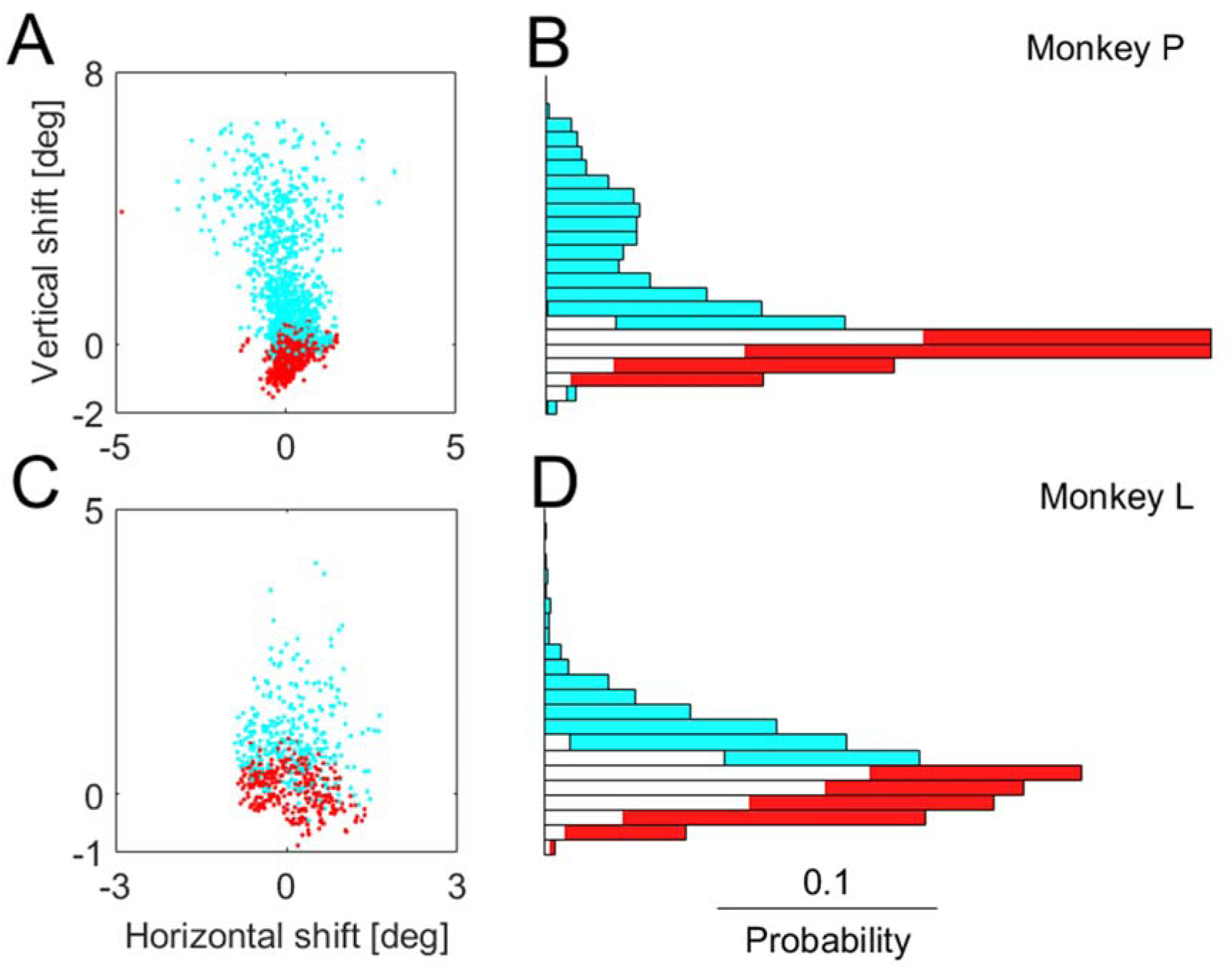
Within-session analysis of the upshift in mesopic vision (cyan) compared to photopic (red). Same format as Fig 5. Statistics included in Table 6.

**Fig 14.**
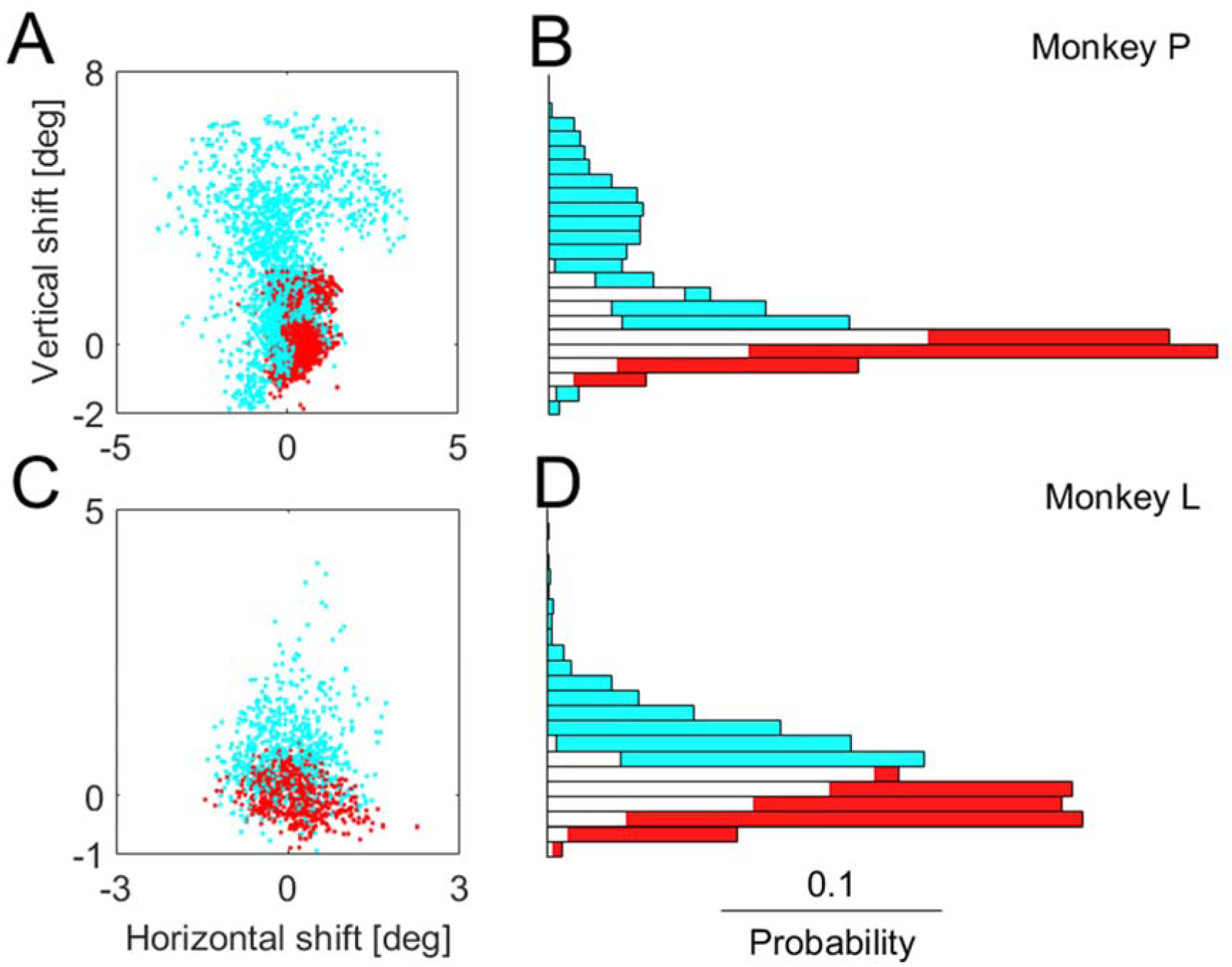
Between-session analysis of the same comparison as Fig 13. Same format as Fig 6. Statistics included in Table 4.

**Table 6.**
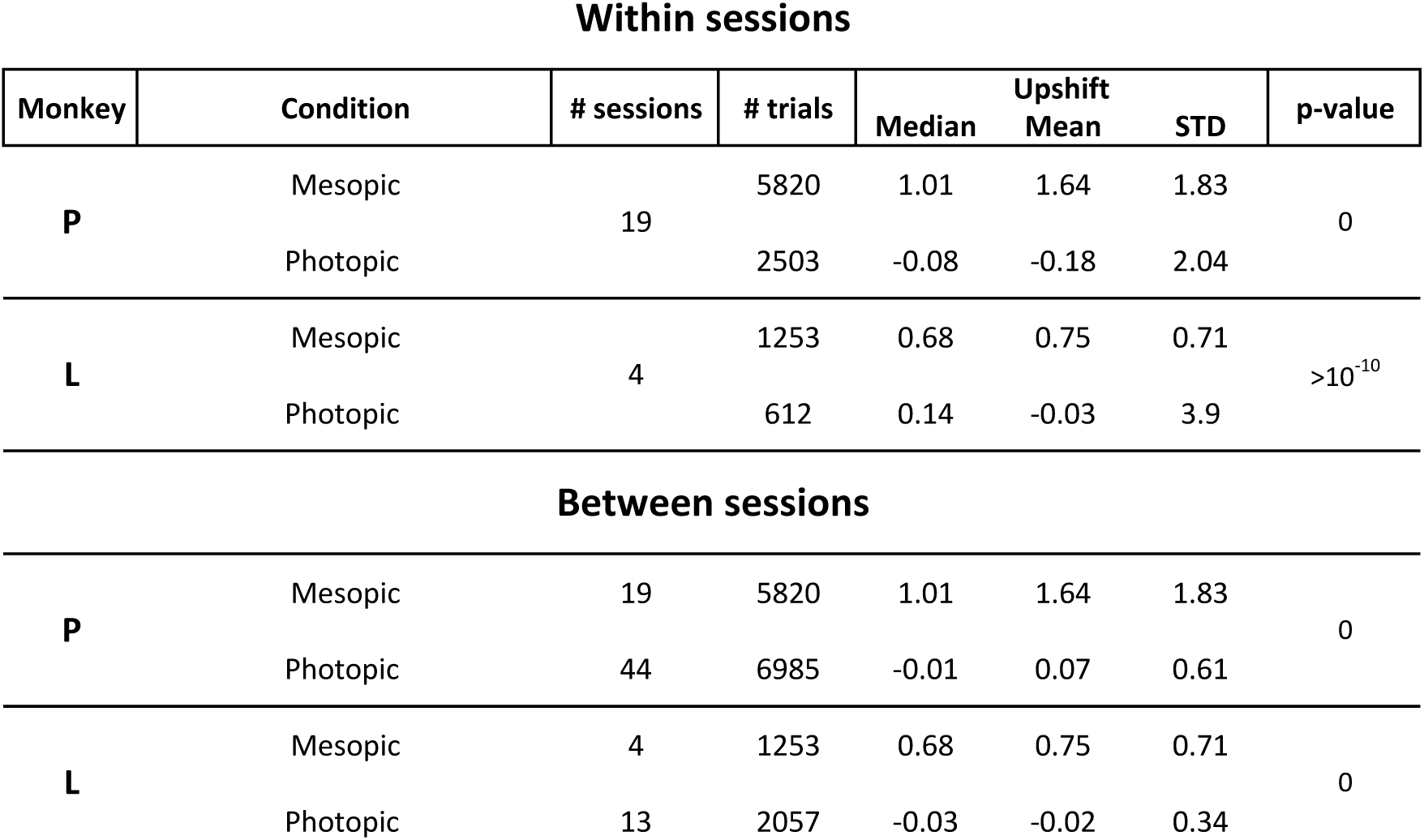
Mesopic vision versus photopic

Within sessions
| Monkey | Condition | # sessions | # trials | Median | Upshift Mean | STD | p-value |
| --- | --- | --- | --- | --- | --- | --- | --- |
| **P** | Mesopic | 19 | 5820 | 1.01 | 1.64 | 1.83 | 0 |
|  | Photopic |  | 2503 | -0.08 | -0.18 | 2.04 |  |
| **L** | Mesopic | 4 | 1253 | 0.68 | 0.75 | 0.71 | >10<sup>-10</sup> |
|  | Photopic |  | 612 | 0.14 | -0.03 | 3.9 |  |
| Between sessions |  |  |  |  |  |  |  |
| **P** | Mesopic | 19 | 5820 | 1.01 | 1.64 | 1.83 | 0 |
|  | Photopic | 44 | 6985 | -0.01 | 0.07 | 0.61 |  |
| **L** | Mesopic | 4 | 1253 | 0.68 | 0.75 | 0.71 | 0 |
|  | Photopic | 13 | 2057 | -0.03 | -0.02 | 0.34 |  |

In both within- and between-sessions comparisons, the cyan dots and histogram bins are above the red ones. The overlap between the red and cyan is not large. As for the null hypothesis, the p-values are less than 10 ^-10^ in all cases (0 in 3 of the 4 comparisons).

**Conclusion:** The null hypothesis is rejected. Mesopic background evokes upshift.

### Mesopic vision induces an intermediate level of upshift

**At issue:** Continuing the previous section, we now aim to test if the upshift is intermediate in value, compared to scotopic-dark background. Recall that the mesopic background follows an interval of dark adaptation, and then testing with scotopic-dark background. Hence, scotopic-dark background is an appropriate comparison for testing if the mesopic upshift is of intermediate values.

**Design:** Condition 1 is mesopic (bright target, mesopic background); condition 2 is scotopic-dark background. Note that the data of condition 1 were always collected after data of condition 2.

The null hypothesis predicts there is no change in upshift between scotopic-dark background and mesopic blocks.

**Results:** Data were collected from 2 monkeys. Fig 15 illustrates the within-session results; Fig 16, the between-sessions. Table 7 specifies the statistics. The Figures follow the pattern described in the previous sections. In both Figures, cyan stands for mesopic trials; blue for scotopic-dark background.

**Fig 15.**
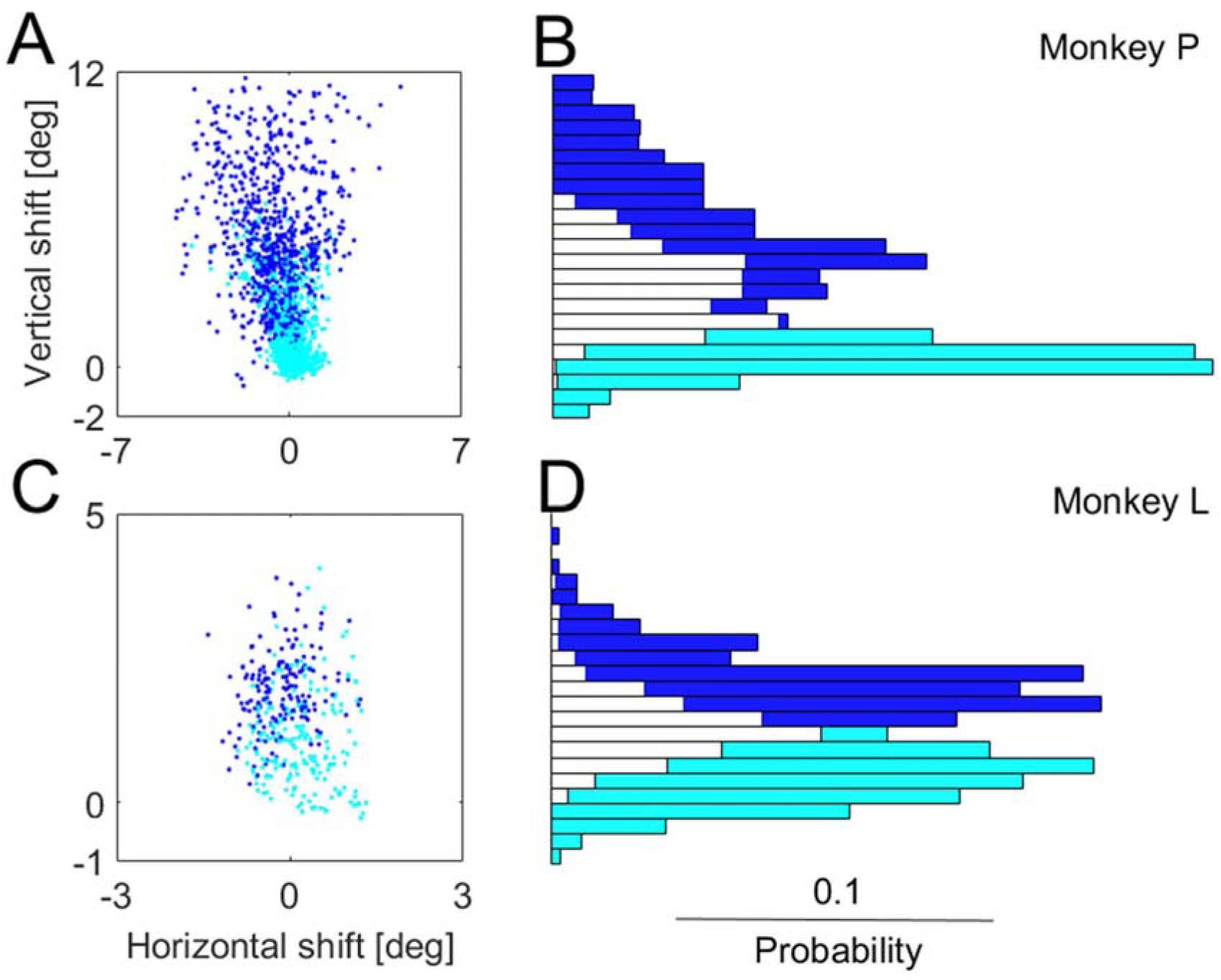
Within-session analysis of the upshift in mesopic vision (cyan) compared to scotopic-dark background (blue). Same format as Fig 5. Statistics included in Table 7.

**Fig 16.**
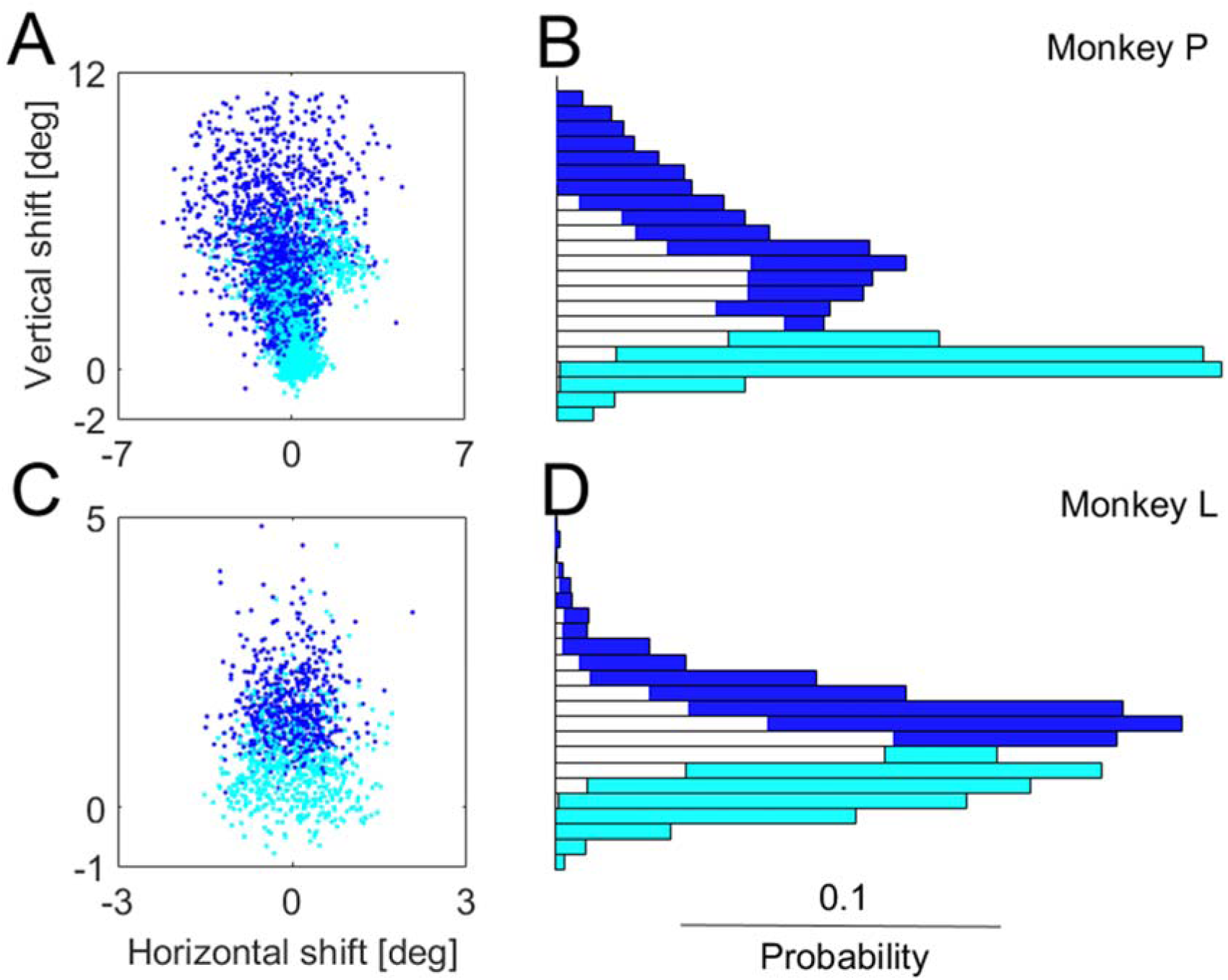
Between-session analysis of the same comparison as Fig 15. Same format as Fig 5. Statistics included in Table 7.

**Table 7.**
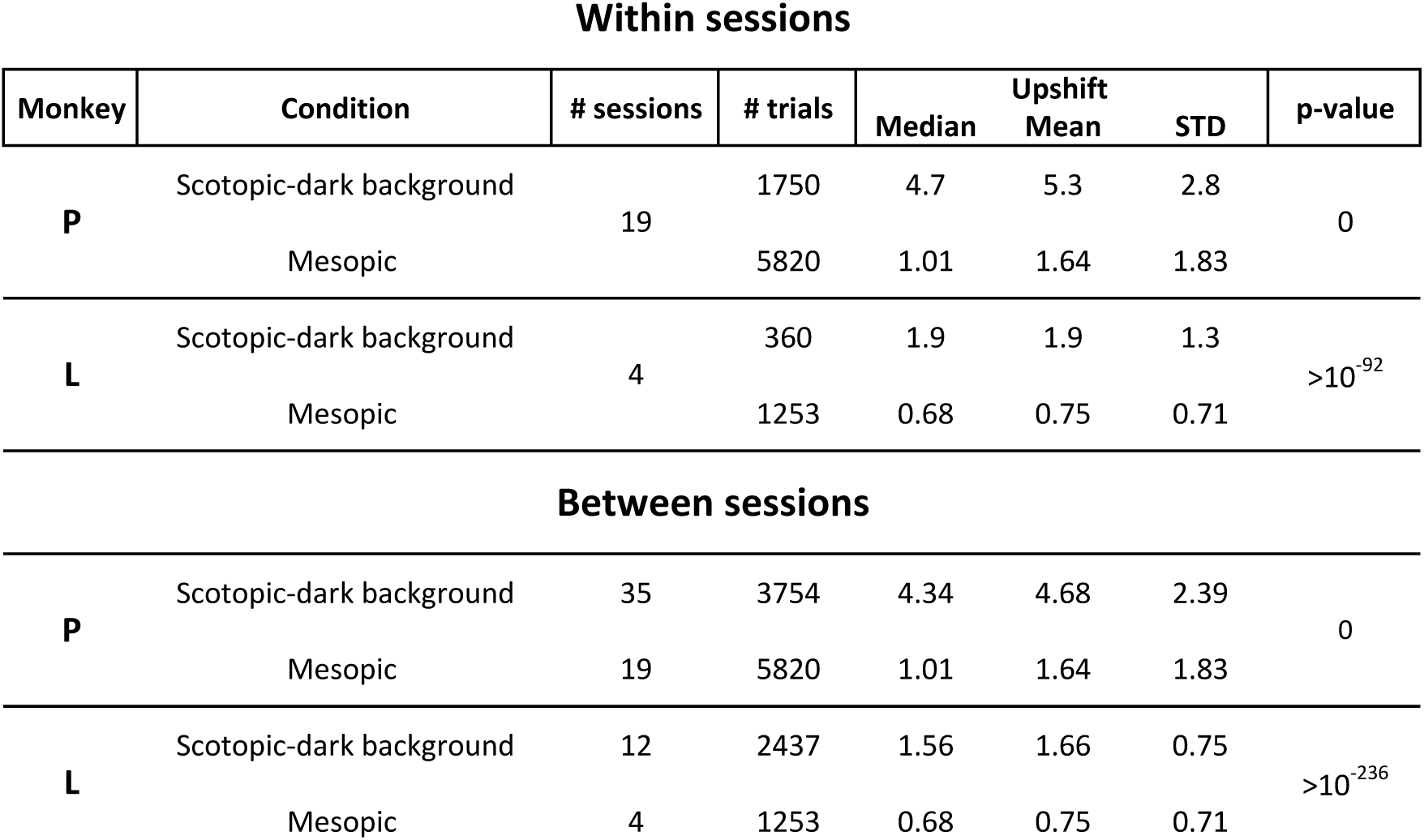
Mesopic vision versus scotopic-dark background

Within sessions
| Monkey | Condition | # sessions | # trials | Median | Upshift Mean | STD | p-value |
| --- | --- | --- | --- | --- | --- | --- | --- |
| **P** | Scotopic-dark background | 19 | 1750 | 4.7 | 5.3 | 2.8 | 0 |
|  | Mesopic |  | 5820 | 1.01 | 1.64 | 1.83 |  |
| **L** | Scotopic-dark background | 4 | 360 | 1.9 | 1.9 | 1.3 | $>10^{-92}$ |
|  | Mesopic |  | 1253 | 0.68 | 0.75 | 0.71 |  |
| Between sessions |  |  |  |  |  |  |  |
| **P** | Scotopic-dark background | 35 | 3754 | 4.34 | 4.68 | 2.39 | 0 |
|  | Mesopic | 19 | 5820 | 1.01 | 1.64 | 1.83 |  |
| **L** | Scotopic-dark background | 12 | 2437 | 1.56 | 1.66 | 0.75 | $>10^{-236}$ |
|  | Mesopic | 4 | 1253 | 0.68 | 0.75 | 0.71 |  |

There is overlap between the histograms in both within and between session analyses. In all cases, the mean of the mesopic upshifts is less than the means of the scotopic-dark background. The null hypothesis is rejected with high significance (p-value less than 10^-92^).

**Conclusion:** Mesopic background evokes upshift, but its size is intermediate, between no upshift (photopic vision) and high upshift (scotopic-dark background).

### No apparent photopic shift on bright adaptation

**At issue:** On going from dark adapted conditions to light, the monkeys’ eyes fixate at lower and lower positions. Can a lengthy exposure to bright light lead to even lower positions than those in the baseline photopic state?

**Design: ‘**Bright adaptation’ consisted of a 45-min waiting interval in bright light. The computer monitor the monkeys faced was set at a level of 7 cd/m^2^, well within the photopic range; more intense background caused discomfort to human observers trying the 45-min interval. In addition, the experimental chamber was illuminated by overhead lights. Condition 1 is standard photopic vision (bright target and background), condition 2 is again photopic vision, but recorded after the bright adaptation interval.

Upshift is computed in the standard fashion. The null hypothesis predicts no systematic change in upshift following bright adaptation.

**Results:** Fig 17 shows the within-session comparison, Fig 18 the between-session. Table 8 shows the statistics.

**Fig 17.**
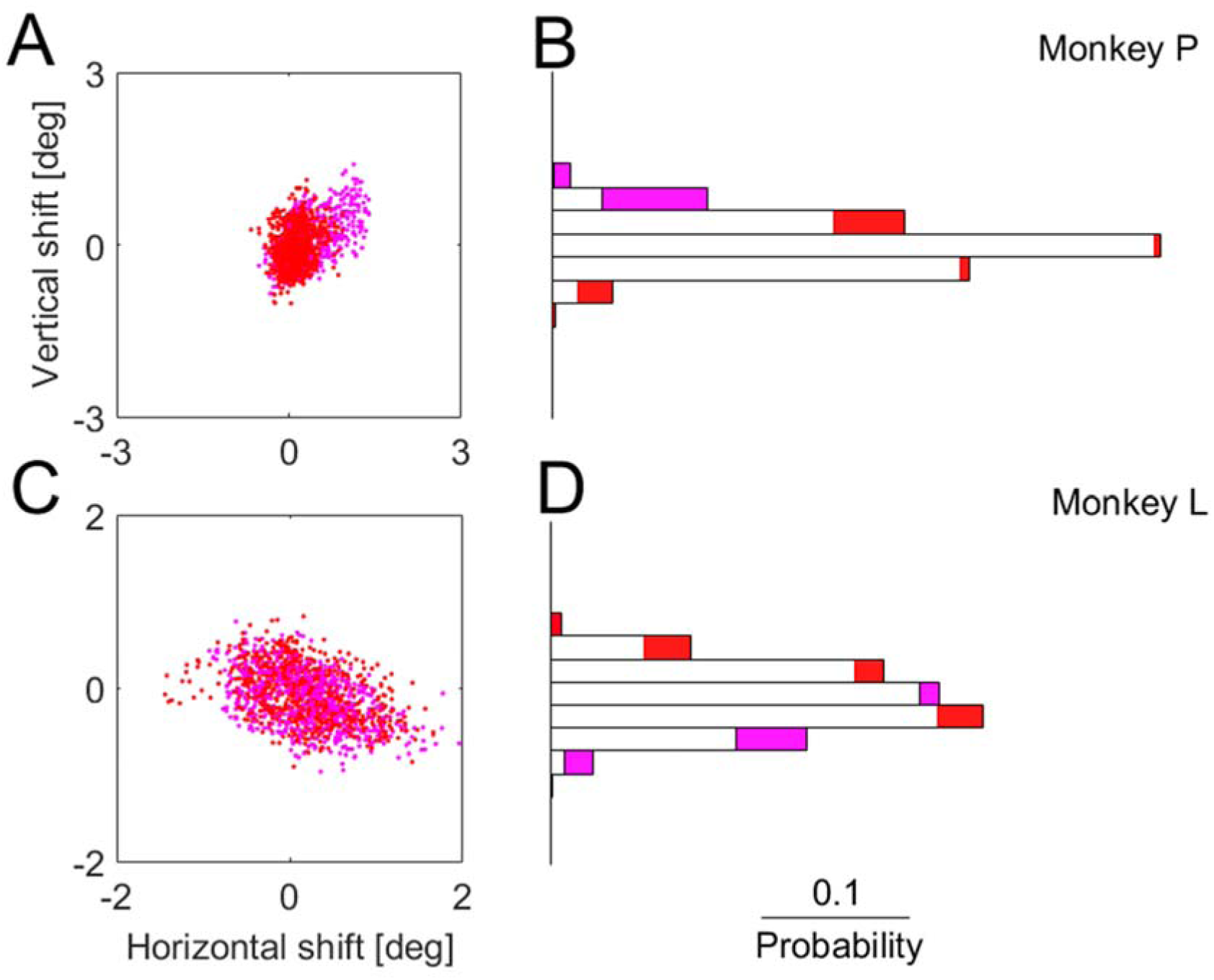
Within-session analysis of the upshift in photopic vision after bright adaptation (pink) compared to baseline photopic (red). Same format as Fig 5. Statistics included in Table 8.

**Fig 18.**
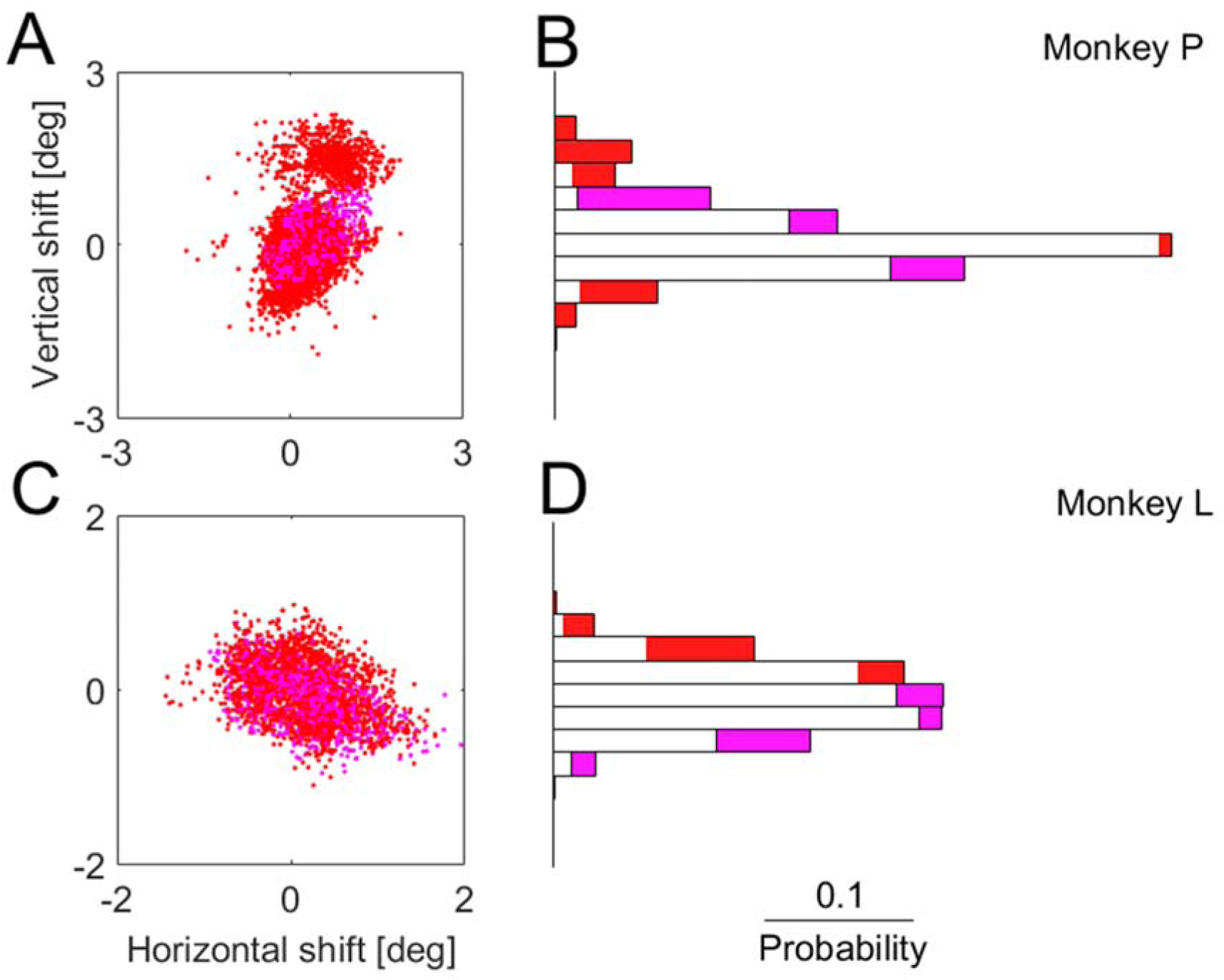
Between-session analysis of the same comparison as Fig 17. Same format as Fig 5. Statistics included in Table 8.

**Table 8.**
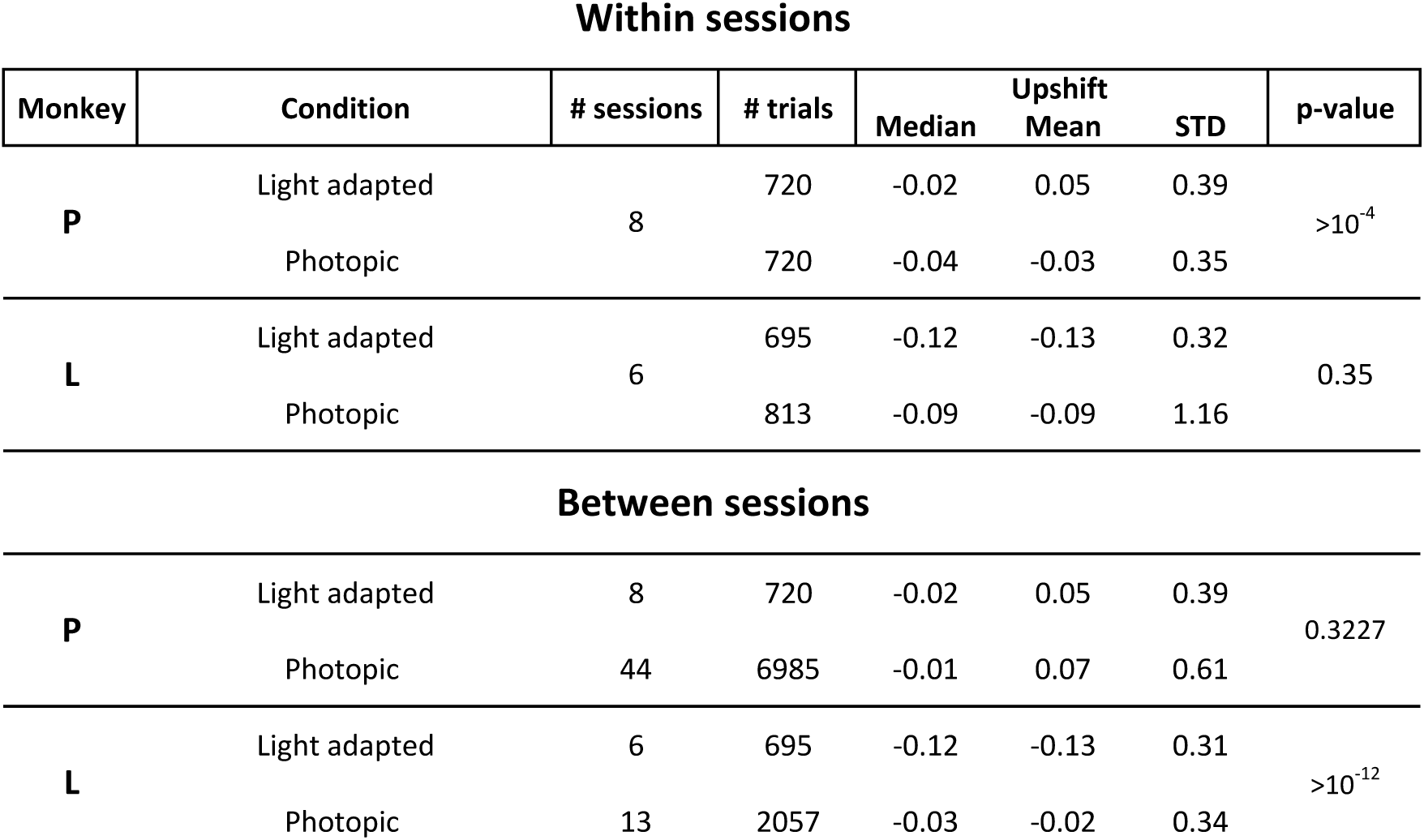
Bright-adapted photopic vision versus baseline photopic vision

Within sessions
| Monkey | Condition | # sessions | # trials | Median | Upshift Mean | STD | p-value |
| --- | --- | --- | --- | --- | --- | --- | --- |
| **P** | Light adapted | 8 | 720 | -0.02 | 0.05 | 0.39 | $>10^{-4}$ |
|  | Photopic |  | 720 | -0.04 | -0.03 | 0.35 |  |
| **L** | Light adapted | 6 | 695 | -0.12 | -0.13 | 0.32 | 0.35 |
|  | Photopic |  | 813 | -0.09 | -0.09 | 1.16 |  |
| **P** | Light adapted | 8 | 720 | -0.02 | 0.05 | 0.39 | 0.3227 |
|  | Photopic | 44 | 6985 | -0.01 | 0.07 | 0.61 |  |
| **L** | Light adapted | 6 | 695 | -0.12 | -0.13 | 0.31 | $>10^{-12}$ |
|  | Photopic | 13 | 2057 | -0.03 | -0.02 | 0.34 |  |

The Within-sessions differences are not significant in Monkey L, and very small in both monkeys – in Monkey P the mean shifts up by 0.08^0^, which is only 23% of the photopic upshift’s standard deviation. The between-session distributions appear more noisy (scatterplots in Fig 18), and Monkey P’s difference is not significant. Monkey L’s mean difference shifts down by 0.090, which is 26% of the photopic standard deviation. Thus, 2 of 4 conditions are not statistically significant, and the other 2 are very small and in opposite directions.

**Conclusion:** Bright adaptation does not bring about upshift or downshift.

## Discussion

### Summary of Results

We studied the eye position with which rhesus monkeys fixate targets in the dark, as compared to bright, photopic illumination. In photopic conditions, the eyes were directed so that the fovea was close to the target. We studied fixation shortly after dark onset, while cones dominate vision (‘photopic dark’), and after 45-min dark adaptation (‘scotopic dark’). In all variations of dark background, the eyes were directed so that the fovea was above the target; thus the image of the target fell on rod-dense superior retina, close to the region of maximal rod density or in between the fovea and that region. The exact target position varied from trial to trial; but the closer to true scotopic the conditions were, the higher was the upshift (bringing the target to higher rod densities). In particular, mesopic vision evoked partial upshift. Dark adaptation (45-min in dark) increased the upshift. There is no analogous photopic effect: 45-min wait in bright, photopic light leads to neither upshift nor downshift.

### The upshift is a continuous parameter

The primary hypothesis leading to the present study is that there are two regions of intensive processing in the retina. In photopic vision, it is the fovea. In scotopic vision, states the hypothesis, it is a rod-dense region, close to those described in studies of monkey retina (Packer et al., 1989; Curcio and Allen, 1990; Wikler and Rakic, 1990; Wikler et al., 1990). Two regions; that would appear to suggest that either, in photopic conditions, the target is fixated with the fovea, hence, no upshift; else, the target is fixated with the rod-dense region in superior retina, hence, upshift. The two anatomical possibilities would seem to translate to two fixation positions. Because the rod-dense region is large and less well-defined than the fovea, some variability could be accepted for the upshifted fixations. However, the variability we observed is much larger. The entire distributions of upshifted fixation positions vary. What is the explanation? Is it consistent with our hypothesis, that the rod-dense region replaces the fovea in scotopic vision?

We suggest that the answer is positive. Apparently in photopic vision fixations always direct the fovea to the target. In full scotopic vision, the upshift has the maximal value; although more variable than photopic fixation, it appears that, by and large, the rod-dense region is used to fixate the target. The higher variability of scotopic fixation positions might have to do with the larger size of the rod-dense region, compared to the fovea.

In intermediate states, intermediate values of upshift are observed. This might be not unlike the saccadic averaging effect: when targets are close, saccades might be directed in between the targets (Findlay, 1982). Somewhat similarly, in situations intermediate between full photopic and full scotopic vision, the eyes might take an intermediate position between the positions of full photopic and full scotopic vision. Thus photopic-dark and scotopic-dark vision both fall in between the extremes of the upshift value, of full photopic and full scotopic vision.

Furthermore, the intermediate values actually taken reflect the visual conditions. Thus, when using a very small bright target, scotopic-dark is functionally closer to scotopic vision than photopic-dark – and, remarkably, the upshift reflects this functional difference.

An additional factor is that in the process of seeing complex objects across saccades, the selection of fixation positions might not be self-evident because there are other processes occurring in parallel (Paeye et al., 2018).

These considerations show that the system’s decision on the exact upshift, the exact fixation position, is not automatic but reflects a subtle assessment of the global visual (and behavioral) conditions.

### The upshift appears to be a trait of scotopic vision

The argument described in the last section shows that the graded values the upshift takes in various behavioral conditions are consistent with the hypothesis of two focal retinal regions, one for photopic vision, the other for scotopic. When behaviorally appropriate, the fixation system chooses an intermediate position between the two centers. Nonetheless, the full phenomenon of upshift occurs in full scotopic vision. Hence upshift is a trait of scotopic vision. It is expressed in full in full scotopic vision. It is expressed in part when conditions call for a combination of photopic and scotopic elements of vision.

### A switch of sensorimotor transformation occurring in nature at least twice a day

Twice a day, it has been known for long, there are transitions occurring between the two visual subsystems, the photopic visual system and the scotopic visual system. For many hours, continuously during daytime, cones mediate vision and at least most rods are just saturated.

Then, after a transition period, rods mediate vision and at least most cones are just too insensitive to be useful. The transition periods, between full scotopic and full photopic states, are rather long themselves.

Now we know that, in rhesus monkeys, together with the sensory switching described above, another switching takes place: switching of the sensorimotor transformations mediating visual fixation. Because the sensory switching is slow, taking more than an hour, the sensorimotor transformation also switches slowly. This might be the ecological setting in which the graded values of upshift have evolved.

Switching of sensorimotor transformations had been studied and one criticism such studies sometimes meet is that they reflect an artificial situation that can be produced in a lab but does not reflect nature. Here we have behaviors that occur spontaneously (recall that the upshift was never rewarded), and they switch on and off in a repeating, fully predictable manner.

## References

Barash S, Melikyan A, Sivakov A, Tauber M. Shift of visual fixation dependent on background illumination. J Neurophysiol 79: 2766–2781, 1998.

Caggiano V, Pomper JK, Fleischer F, Fogassi L, Giese M, Thier P. Mirror neurons in monkey area F5 do not adapt to the observation of repeated actions. Nat Commun 4: 1433, 2013.

Curcio CA, Allen KA. Topography of ganglion cells in human retina. J Comp Neurol 300: 5–25, 1990.

Dash S, Catz N, Dicke PW, Thier P. Encoding of smooth-pursuit eye movement initiation by a population of vermal Purkinje cells. Cereb Cortex 22: 877–891, 2012.

Findlay JM. Global visual processing for saccadic eye movements. Vision Res 22: 1033–1045, 1982.

Normann R, Werblin F. CONTROL OF RETINAL SENSITIVITY.1. LIGHT AND DARK-ADAPTATION OF VERTEBRATE RODS AND CONES. 63: 37–61, 1974.

Packer O, Hendrickson AE, Curcio CA. Photoreceptor topography of the retina in the adult pigtail macaque (Macaca nemestrina). J Comp Neurol 288: 165–183, 1989.

Paeye C, Collins T, Cavanagh P, Herwig A. Calibration of peripheral perception of shape with and without saccadic eye movements. Atten Percept Psychophys 80: 723–737, 2018.

Spivak O, Thier P, Barash S. Persistence of the dark-background-contingent gaze upshift during visual fixations of rhesus monkeys. J Neurophysiol 112: 1999–2005, 2014.

Wikler KC, Rakic P. Distribution of photoreceptor subtypes in the retina of diurnal and nocturnal primates. J Neurosci 10: 3390–3401, 1990.

Wikler KC, Williams RW, Rakic P. Photoreceptor mosaic: number and distribution of rods and cones in the rhesus monkey retina. J Comp Neurol 297: 499–508, 1990.

